# A *Drosophila* screen of schizophrenia-related genes highlights the requirement of neural and glial matrix metalloproteinases for neuronal remodeling

**DOI:** 10.1101/2024.10.01.616025

**Authors:** Shir Keret, Hagar Meltzer, Neta Marmor-Kollet, Oren Schuldiner

## Abstract

Schizophrenia (SCZ) is a multifactorial neuropsychiatric disorder of complex and mostly unknown etiology, affected by genetic, developmental and environmental factors. Neuroanatomical abnormalities, such as loss of grey matter, are apparent prior to the onset of symptoms, suggesting neurodevelopmental origin. Indeed, it has been hypothesized, and recently experimentally supported, that SCZ is associated with dysregulation of developmental synaptic pruning. Here, we explore the molecular link between SCZ-associated genes and developmental neuronal remodeling. We focused on the *Drosophila* mushroom body (MB), which undergoes stereotypic remodeling during metamorphosis. We conducted a loss-of-function screen in which we knocked down, in either glia or neurons, *Drosophila* homologs of human genes that are associated with SCZ based on genomic studies (GWAS). Out of our ‘positive hits’, we focused on matrix metalloproteinases (MMPs), mostly known for their role in remodeling of the extracellular matrix. Our combinatorial loss-of-function experiments suggest that *Drosophila* MMPs, which are closely related to mammalian MMPs, are required in neurons and in glia for the pruning of MB axons. Our results shed new light on potential molecular players underlying neurodevelopmental defects in SCZ and highlight the advantage of genetically tractable model organisms in the study of human neurodevelopmental disorders.

## Introduction

Following its initial establishment, the developing nervous system of both vertebrates and invertebrates undergoes remodeling to shape its mature connectivity. In humans, developmental neuronal remodeling occurs postnatally and includes degenerative events such as retraction of synapses and large-scale axon elimination. While these processes occur most prominently during the first two years of life, they continue until after adolescence (Luo & O’Leary, 2005; Schuldiner & Yaron, 2015). Defects in neuronal remodeling have long been hypothesized to be one of the underlying causes of neuropsychiatric disorders, including schizophrenia (SCZ; Feinberg, 1982; Penzes et al., 2011).

SCZ is a severe, multifactorial mental health condition affecting ∼1% of the population. It is characterized by delusions, hallucinations, social withdrawal and cognitive deficits, typically onsetting during the late teens to early adulthood (McCutcheon et al., 2020). It is known from MRI studies that structural abnormalities in the brain, such as reduction in gray matter and ventricular enlargement, may precede the onset of psychotic symptoms, implying neurodevelopmental origin (Brent et al., 2013; Omlor et al., 2025; Suzuki et al., 2002; Zikidi et al., 2020). It has been suggested, for many years, that the characteristic neuroanatomical alterations in individuals with SCZ are due to dysregulation of neuronal remodeling, specifically over-pruning of synapses (Feinberg, 1982; Hoffman & Dobscha, 1989; Huttenlocher & Dabholkar, 1997; Keshavan et al., 1994; Peter R, 1979). The first experimental evidence to support this hypothesis came several years ago, and associated SCZ with excessive expression of complement components leading to increased synapse elimination by microglia (Sekar et al., 2016). This work as well as others highlighted glia, including microglia and astrocytes, as key mediators of synapse pruning in normal development as well as in SCZ and other neuropathologies (Reviewed in Laricchiuta et al., 2024; Notter, 2021; Scott-Hewitt et al., 2023). Still, the molecular basis of SCZ risk and its link to abnormalities in neuronal remodeling remains mostly unknown.

*Drosophila melanogaster* is an ideal model to study neuronal remodeling, due to its unparalleled genetic toolkit combined with its massive stereotypic circuit remodeling during metamorphosis (Truman, 1990; Yaniv & Schuldiner, 2016). Our focus is on the *Drosophila* mushroom body (MB), a brain structure that functions as a center of olfactory learning and memory (Heisenberg, 1998). The MB is comprised of three types of intrinsic neurons, known as Kenyon cells (KCs), which are sequentially born from identical neuroblasts (Fig. 1A). Out of the three KC types (γ, ɑ’/β’ and ɑ/β), only the first-born γ- KCs undergo stereotypic remodeling. When they initially grow, in the embryonic and early larval stages, γ-KCs project axons that bifurcate to form vertical and medial lobes. At the early pupal stage (∼6-18 hours after puparium formation; h APF) both axonal lobes prune up to their branchpoint. Subsequently, γ-KCs initiate axon regrowth to create the adult-specific medially projecting γ-lobe (Lee et al., 1999; Yaniv & Schuldiner, 2016; Fig. 1A). Importantly, glia were shown, by our lab and others, to actively participate in the remodeling of γ-KCs (Awasaki et al., 2011; Awasaki & Ito, 2004; Boulanger et al., 2021; Hakim et al., 2014; Marmor-Kollet et al., 2023; Watts et al., 2004). Three glial subtypes are present around the MB: cortex glia, ensheathing glia and astrocyte-like glia (hereafter referred to as astrocytes; Freeman, 2015). Astrocytes are the main scavengers of axonal debris following pruning (Hakim et al., 2014; Tasdemir-Yilmaz & Freeman, 2014). Additionally, inhibiting astrocytic functions results in brain-wide defects in synapse elimination (Tasdemir-Yilmaz & Freeman, 2014). Moreover, we recently demonstrated that astrocytes actively infiltrate the axon bundle prior to pruning to facilitate axon defasciculation and elimination (Marmor-Kollet et al., 2023).

**Figure 1.**
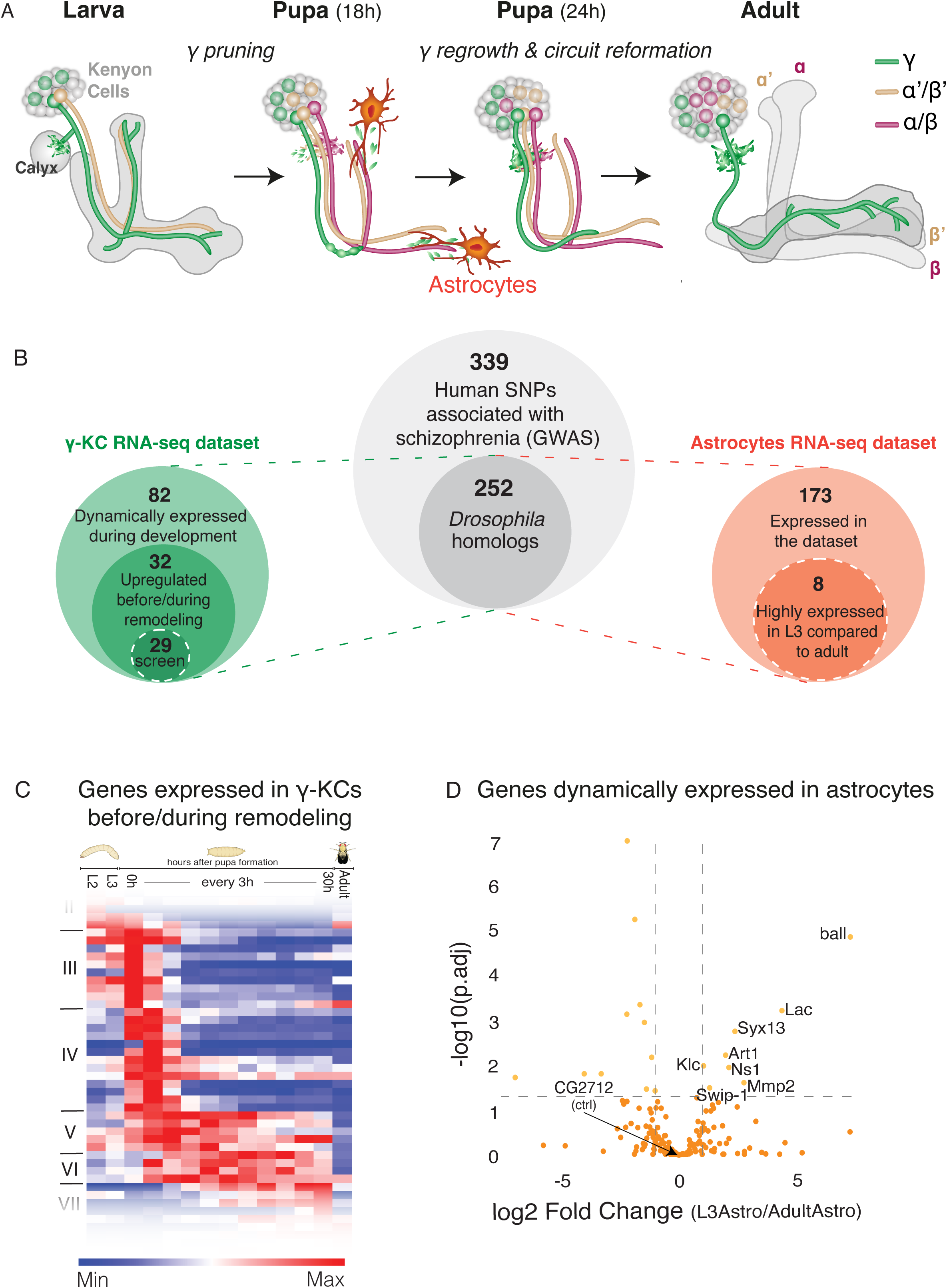
*Drosophila* homologs of SCZ-associated genes are dynamically expressed in developing mushroom body γ-KCs and in astrocytes. (A) Schematic representation of *Drosophila* MB development. During early pupal stages, only the γ-KCs (green) prune their dendrites completely and their vertical and medial axonal branches up to their branch-point. Subsequently, they reform dendrites in the calyx and regrow axons to form the adult-specific medial lobe. Later-born α’/β’- and α/β-KCs, as well as astrocytes, are also depicted. (B) A schematic representation describing the logic of candidate selection for the loss-of-function screen within KCs and glia. White dashed circles highlight the portion of genes that were screened. (C) A heatmap representing the expression of 32 genes (clusters III-VI) upregulated during/after remodeling that were selected as candidates for screening (based on Alyagor et al., 2018). Each row represents a gene, with red and blue indicating high and low relative expression, respectively. The full heatmap of all dynamically expressed genes is available in Fig. S1. (D) Volcano plot representing the comparison of gene expression between larval and adult astrocytes (based on Marmor-Kollet et al., 2023). Annotated genes are expressed significantly higher in L3 astrocytes compared to adult astrocytes. Each dot represents a gene, the x axis represents the log2 fold change, the y axis represents the −log10 of the adjusted p value (p.adj). For the full gene expression data and the candidate gene lists see Supplemental excel file 1.

*Drosophila* is routinely used to study human neurodegenerative and neuropsychiatric disorders (Lessing & Bonini, 2009; van Alphen & van Swinderen, 2013), and SCZ-associated genes were shown to affect MB structure, activity and function in memory and sleep (Furukubo-Tokunaga, 2009; Hidalgo et al., 2021; Sawamura et al., 2008). Here, we harness the fly genetic toolkit and the well-characterized remodeling of the MB to delve into the molecular underpinnings of the long-standing hypothesis that defects in neuronal remodeling contribute to SCZ etiology. We systematically examine how MB remodeling is affected by perturbations in *Drosophila* genes whose human homologs have been associated with SCZ. Our findings identify novel players in axon pruning and further highlight the significance of glial contribution to remodeling, providing a solid basis for future research on the neurodevelopmental molecular origin of SCZ.

## Results

### Drosophila homologs of human SCZ-associated genes are dynamically expressed in developing mushroom body γ-KCs and in astrocytes

We searched genome-wide association studies (GWAS) for candidate genes that have single nucleotide polymorphism (SNPs) in SCZ patients (339 genes; Hamshere et al., 2013; Wu et al., 2017; szdb.org/index.html), which is a compilation of data from the Psychiatric Genomics Consortium (PGC) and CLOZUK. We converted the SNP-containing genes to their *Drosophila* homologs using HumanMine (Lyne et al., 2022), resulting in 252 genes (Fig. 1B; Supplementary excel file 1). To narrow our candidate list to genes that are potentially required for MB pruning, we first used our previously generated transcriptional atlas of developing γ-KCs (Alyagor et al., 2018) to identify 196 genes that are expressed above threshold levels, out of which 82 showed dynamic expression (Fig. S1). Next, we narrowed our list to genes that are specifically upregulated during or prior to γ-axon pruning (Fig. 1B, Fig. 1C clusters III-VI highlighted). This process resulted in 32 candidate genes, out of which 29 had available RNA-interference (RNAi) fly lines for loss-of-function (LOF) screening (Fig.1B; Supplementary excel file 1). Due to the known significance of glia in pruning, in parallel we also examined a second transcriptional dataset, of larval and adult astrocytes, previously generated in the lab (Marmor-Kollet et al., 2023). Out of the 252 *Drosophila* homologs of SCZ-associated genes, 173 are expressed in astrocytes above threshold levels (Fig.1B; Supplementary excel file 1). Out of these, 8 are upregulated in late larval (3^rd^ instar larvae; L3) compared to adult astrocytes (Fig. 1D; Supplementary excel file 1), suggesting a potential role in axon pruning. All 8 genes had available RNAi lines for screening.

### Drosophila homologs of human SCZ-associated genes are required in neurons and glia for γ-axon pruning

We screened candidate genes using a LOF strategy by tissue-specific expression of RNAi-species via the Gal4-UAS system (Brand & Perrimon, 1993). For KD in γ-KCs, we used the pan-KC driver OK107-Gal4, which is strongly expressed in all KC types (ɑ/β, ɑ’/β’ and γ), combined with a second, independent binary system (Riabinina & Potter, 2016) to specifically visualize γ-KCs in the adult MB (R71G10-QF2 driving mtdT-HA; Fig. 2A). Notably, although remodeling of MB γ-KCs occurs during metamorphosis, unpruned larval axons persist until adulthood. Thus, adult MBs reflect abnormalities that occurred during development. Out of the 29 genes we screened, OK107-Gal4-driven KD of 4 genes resulted in lethality, 11 genes displayed varying degrees of pruning defects (‘positive hits’; Fig. 2B-N), and one gene led to abnormal MB morphology (Fig. S2). In parallel, we used a similar strategy to screen the 8 candidate genes in glia - by driving RNAi using the strong pan-glial driver Repo-Gal4, combined with R71G10-QF2-driven mtdT to visualize γ-KCs (Fig. 2O). Strikingly, KD of 7 of the 8 genes resulted in γ-axon pruning defects (Fig. 2P-X).

**Figure 2.**
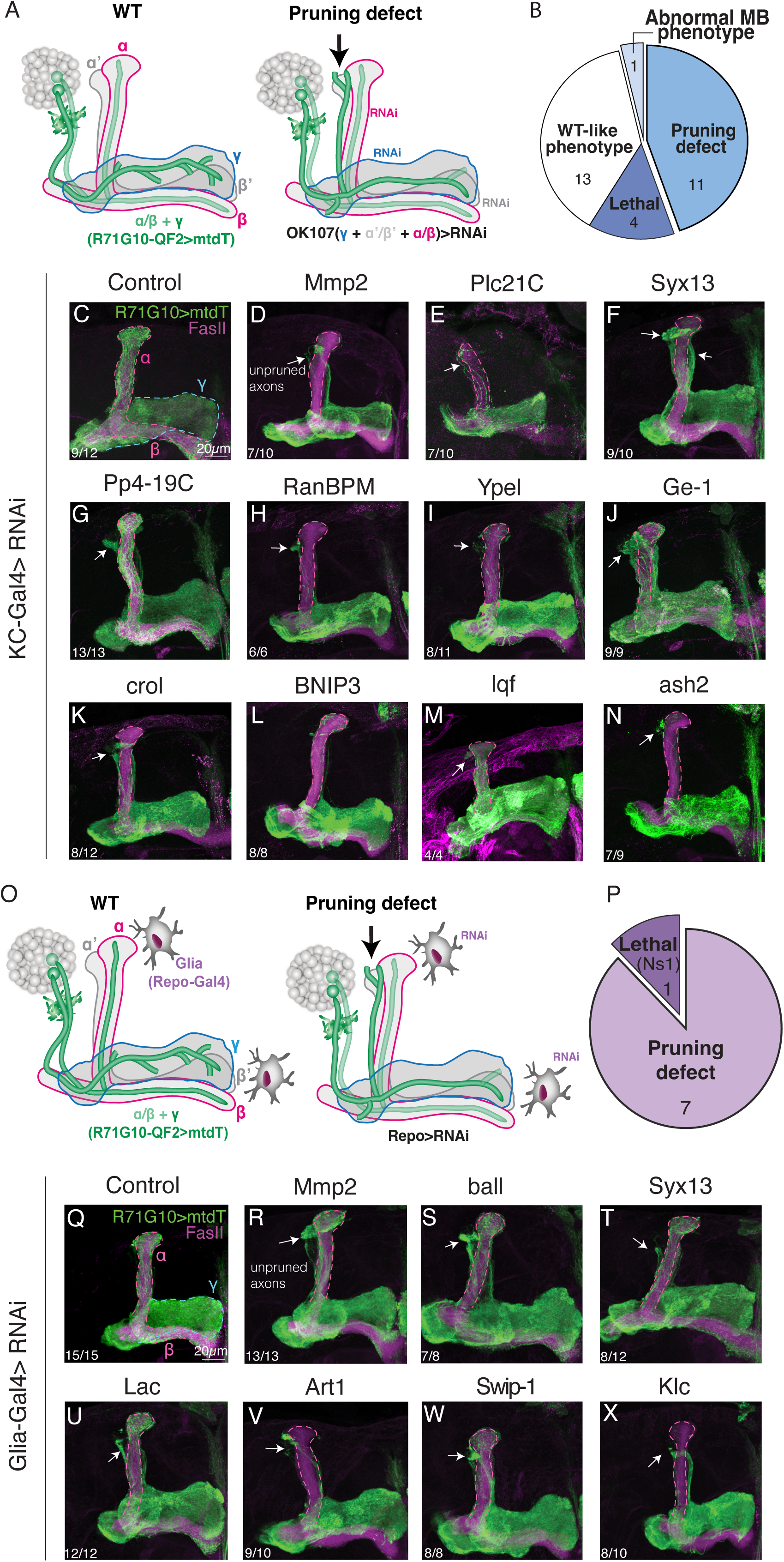
*Drosophila* homologs of human SCZ-associated genes are required in neurons and glia for γ-axon pruning. (A) A schematic representation of the experimental design. RNAi transgenes targeting candidate genes were driven in KCs by the strong driver OK107-Gal4 (γ - blue outline, α/β - magenta outline, α’/β’ - gray). The γ-axon pruning phenotype was examined in adult brains via the expression of myristoylated tandem Tomato (mtdT; green) driven by R71G10-QF2. Left: WT, all γ axons were pruned during pupal stages and regrew to form the adult specific medial γ lobe (blue outline). Note that R71G10-QF2 is also stochastically expressed in ⍺/β neurons, which are additionally strongly positive to FasII (magenta outline). Right: pruning defect, some γ axons were not pruned - remanents of larval projections (arrow) are visible outside the highly fasciculated α axon bundle (magenta outline). Adapted with permission from Marmor-Kollet et al. 2023. (B) Pie chart representing the overall KD screen results in KCs (see Supplemental excel file 1; for the abnormal MB phenotype see Fig. S2). (C-N) Confocal z-projections of of adult MBs in which the indicated UAS-RNAi transgenes were expressed in all KCs using OK107-Gal4, while R71G10-QF2 drives the expression of mtdT (green) in γ-KCs and stochastically in ɑ/β-KCs. The RNAi transgenes target the following genes: Matrix Metalloproteinase 2 (Mmp2; D), Phospholipase C at 21C (Plc21C; E), Syntaxin 13 (Syx13; F), Protein phosphatase 19C (Pp4-19C; G), Ran-binding protein M (RanBPM; H), Yippee-like (Ypel; I), Ge-1 (Ge-1; J), crooked legs (crol; K), BCL2 Interacting Protein 3 (BNIP3; L), liquid facets (lqf; M), absent, small, or homeotic discs 2 (ash2; N). Control is RNAi targeting Luciferase (C). FasII antibody (magenta) strongly labels α/β-KCs and weakly labels γ-KCs. The γ lobe is outlined in blue, and the α/β lobes are outlined in magenta. (O) A schematic representation of the experimental design. RNAi transgenes targeting candidate genes were driven in all glia by Repo-Gal4 (gray cells). The γ axon pruning phenotype was examined in adult brains via the expression of mtdT (green) driven by R71G10-QF2. (P) Pie chart representing the overall glial screen results (see Supplemental excel file 1). (Q-X) Confocal z-projections of adult MBs in which the indicated UAS-RNAi transgenes were expressed in all glia cells using Repo-Gal4, while R71G10-QF2-drives the expression of mtdT (green) in γ-KCs and stochastically in ɑ/β-KCs. The RNAi transgenes target the following genes: Matrix Metalloproteinase 2 (Mmp2; R), ballchen (ball; S), Syntaxin 13 (Syx13; T), Lachesin (Lac; U), Arginine methyltransferase 1 (Art1; V), Swiprosin-1 (Swip-1; W), Kinesin light chain (Klc; X). Control is RNAi targeting CG2712 (Q), which is expressed in near-zero levels (in both larval and adult astrocytes), as indicated by its location at the origin of the volcano plot (see Fig. 1D). FasII antibody (magenta) strongly labels α/β-axons and weakly labels γ-axons. The γ lobe is outlined in blue, and the α/β lobes are outlined in magenta. Arrows indicate unpruned axons. Scale bar corresponds to 20µm. The number of MBs (each from an individual brain) showing the presented phenotype out of the total n for each genotype is indicated. For all screened genes genotype and phenotype findings, see Supplemental excel file 1. Lines used in the screen appear in Table 1. Full genotypes appear in Table 2.

**Table 1.**
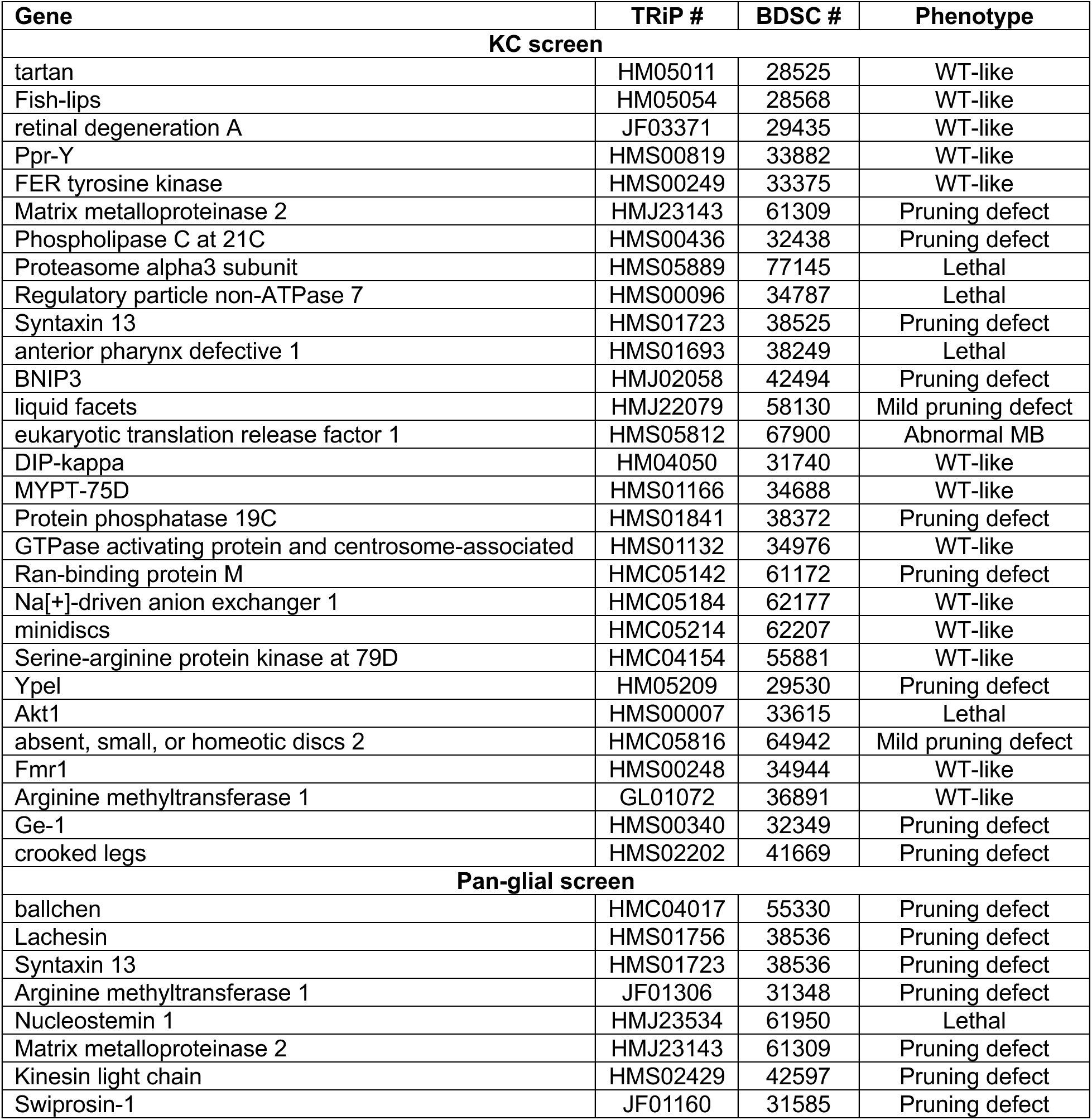
RNAi lines used in the KC screen (using OK107-Gal4) and the pan-glial screen (using Repo-Gal4) and their KD phenotypes.

**Table 2.**
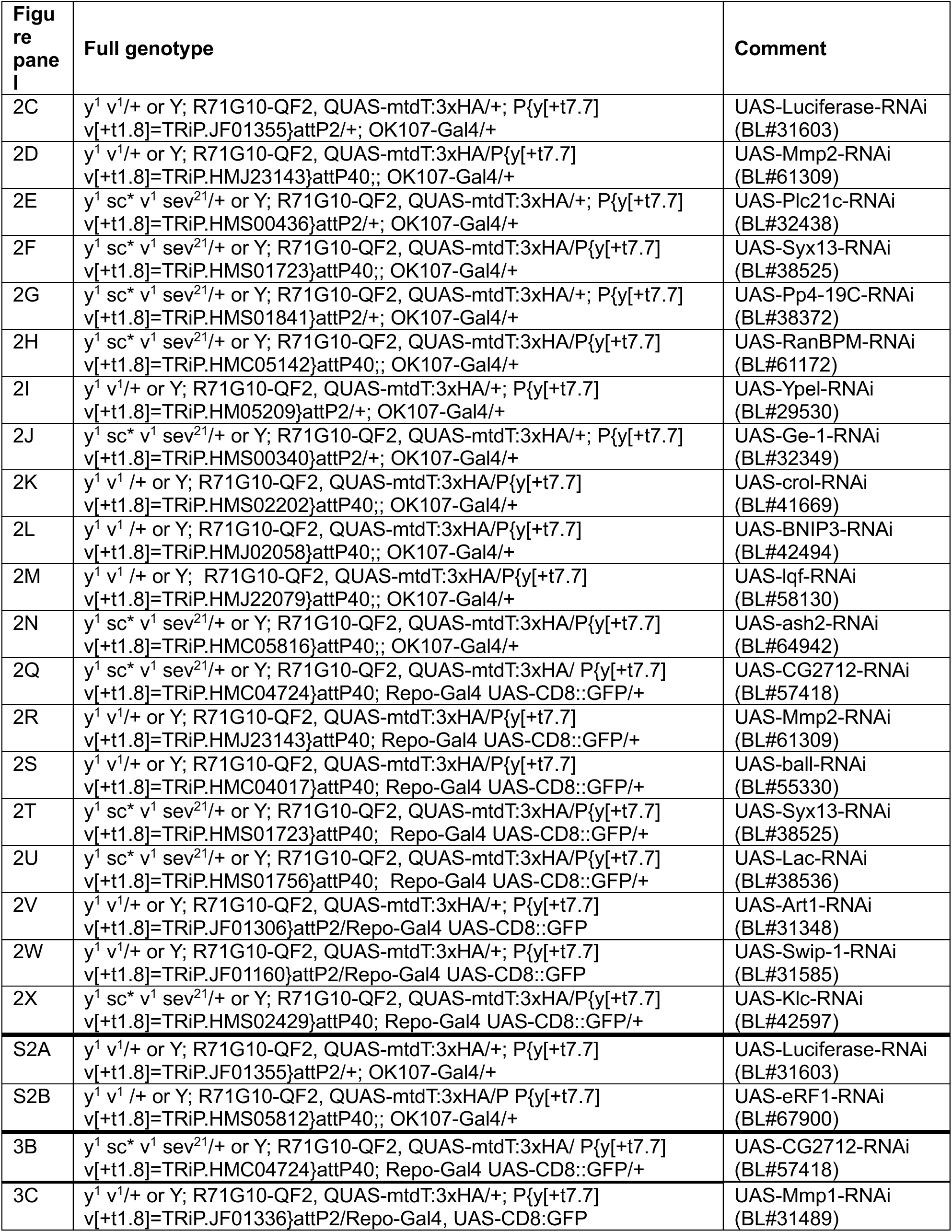
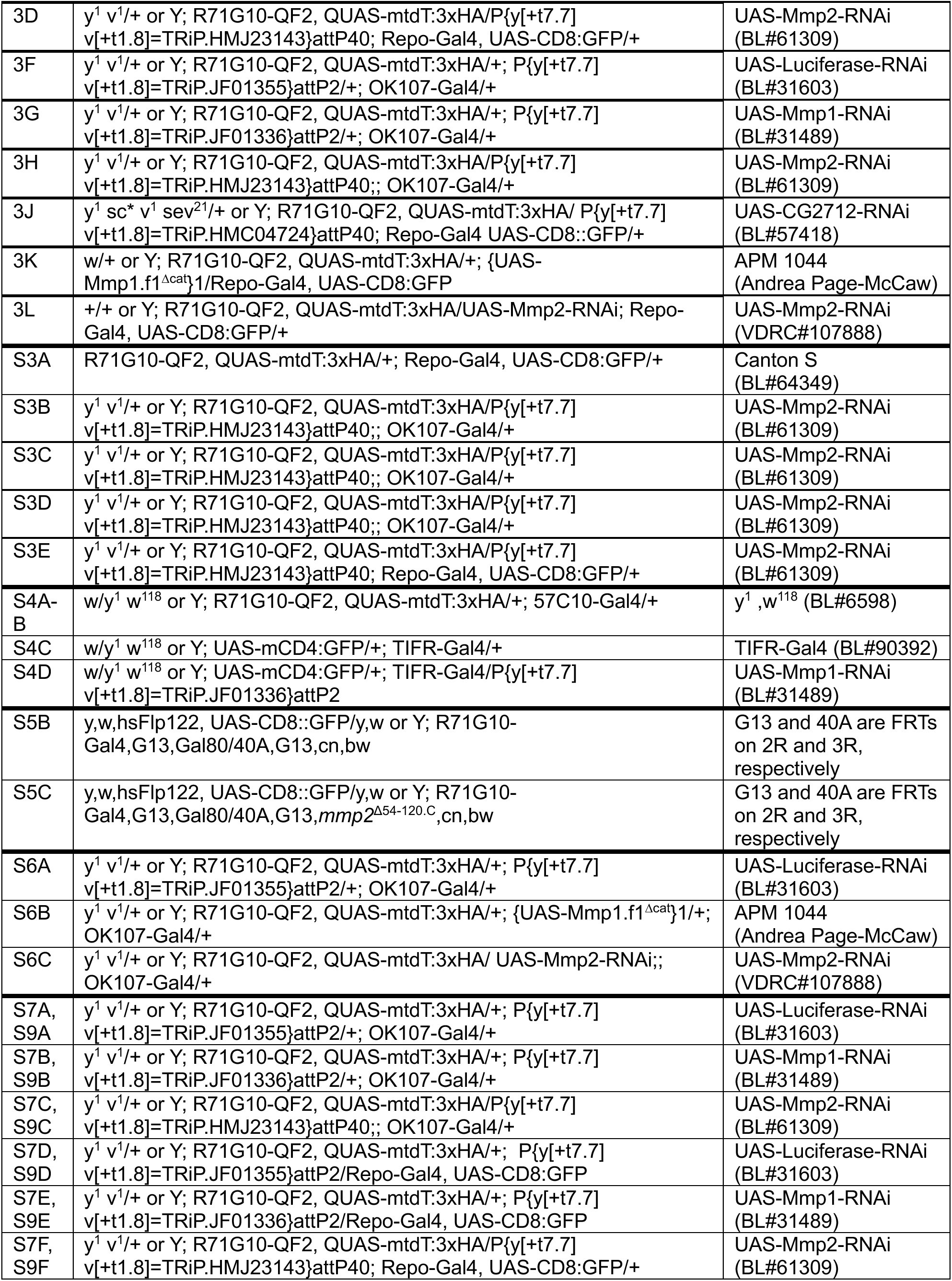
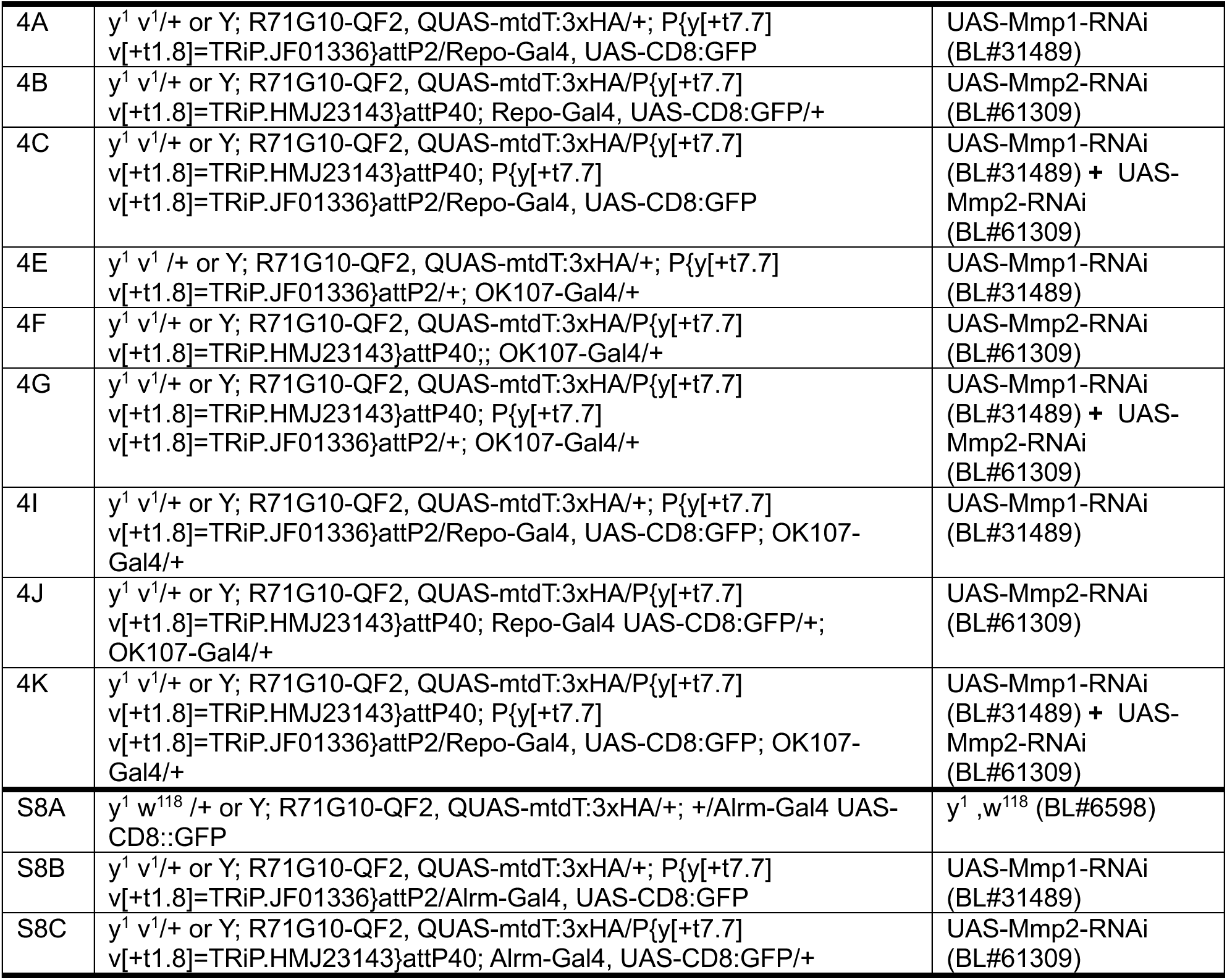
Full *Drosophila* genotypes.

| Figure panel | Full genotype | Comment |
| --- | --- | --- |
| 2C | y<sup>1</sup> v<sup>1</sup>/+ or Y; R71G10-QF2, QUAS-mtdT:3xHA/+; P{y[+t7.7] v[+t1.8]=TRiP.JF01355}attP2/+; OK107-Gal4/+ | UAS-Luciferase-RNAi (BL#31603) |
| 2D | y<sup>1</sup> v<sup>1</sup>/+ or Y; R71G10-QF2, QUAS-mtdT:3xHA/P{y[+t7.7] v[+t1.8]=TRiP.HMJ23143}attP40;; OK107-Gal4/+ | UAS-Mmp2-RNAi (BL#61309) |
| 2E | y<sup>1</sup> sc* v<sup>1</sup> sev<sup>21</sup>/+ or Y; R71G10-QF2, QUAS-mtdT:3xHA/+; P{y[+t7.7] v[+t1.8]=TRiP.HMS00436}attP2/+; OK107-Gal4/+ | UAS-Plc21c-RNAi (BL#32438) |
| 2F | y<sup>1</sup> sc* v<sup>1</sup> sev<sup>21</sup>/+ or Y; R71G10-QF2, QUAS-mtdT:3xHA/P{y[+t7.7] v[+t1.8]=TRiP.HMS01723}attP40;; OK107-Gal4/+ | UAS-Syx13-RNAi (BL#38525) |
| 2G | y<sup>1</sup> sc* v<sup>1</sup> sev<sup>21</sup>/+ or Y; R71G10-QF2, QUAS-mtdT:3xHA/+; P{y[+t7.7] v[+t1.8]=TRiP.HMS01841}attP2/+; OK107-Gal4/+ | UAS-Pp4-19C-RNAi (BL#38372) |
| 2H | y<sup>1</sup> sc* v<sup>1</sup> sev<sup>21</sup>/+ or Y; R71G10-QF2, QUAS-mtdT:3xHA/P{y[+t7.7] v[+t1.8]=TRiP.HMC05142}attP40;; OK107-Gal4/+ | UAS-RanBPM-RNAi (BL#61172) |
| 2I | y<sup>1</sup> v<sup>1</sup>/+ or Y; R71G10-QF2, QUAS-mtdT:3xHA/+; P{y[+t7.7] v[+t1.8]=TRiP.HM05209}attP2/+; OK107-Gal4/+ | UAS-Ypel-RNAi (BL#29530) |
| 2J | y<sup>1</sup> sc* v<sup>1</sup> sev<sup>21</sup>/+ or Y; R71G10-QF2, QUAS-mtdT:3xHA/+; P{y[+t7.7] v[+t1.8]=TRiP.HMS00340}attP2/+; OK107-Gal4/+ | UAS-Ge-1-RNAi (BL#32349) |
| 2K | y<sup>1</sup> v<sup>1</sup> /+ or Y; R71G10-QF2, QUAS-mtdT:3xHA/P{y[+t7.7] v[+t1.8]=TRiP.HMS02202}attP40;; OK107-Gal4/+ | UAS-crol-RNAi (BL#41669) |
| 2L | y<sup>1</sup> v<sup>1</sup> /+ or Y; R71G10-QF2, QUAS-mtdT:3xHA/P{y[+t7.7] v[+t1.8]=TRiP.HMJ02058}attP40;; OK107-Gal4/+ | UAS-BNIP3-RNAi (BL#42494) |
| 2M | y<sup>1</sup> v<sup>1</sup> /+ or Y; R71G10-QF2, QUAS-mtdT:3xHA/P{y[+t7.7] v[+t1.8]=TRiP.HMJ22079}attP40;; OK107-Gal4/+ | UAS-lqf-RNAi (BL#58130) |
| 2N | y<sup>1</sup> sc* v<sup>1</sup> sev<sup>21</sup>/+ or Y; R71G10-QF2, QUAS-mtdT:3xHA/P{y[+t7.7] v[+t1.8]=TRiP.HMC05816}attP40;; OK107-Gal4/+ | UAS-ash2-RNAi (BL#64942) |
| 2Q | y<sup>1</sup> sc* v<sup>1</sup> sev<sup>21</sup>/+ or Y; R71G10-QF2, QUAS-mtdT:3xHA/ P{y[+t7.7] v[+t1.8]=TRiP.HMC04724}attP40; Repo-Gal4 UAS-CD8::GFP/+ | UAS-CG2712-RNAi (BL#57418) |
| 2R | y<sup>1</sup> v<sup>1</sup>/+ or Y; R71G10-QF2, QUAS-mtdT:3xHA/P{y[+t7.7] v[+t1.8]=TRiP.HMJ23143}attP40; Repo-Gal4 UAS-CD8::GFP/+ | UAS-Mmp2-RNAi (BL#61309) |
| 2S | y<sup>1</sup> v<sup>1</sup>/+ or Y; R71G10-QF2, QUAS-mtdT:3xHA/P{y[+t7.7] v[+t1.8]=TRiP.HMC04017}attP40; Repo-Gal4 UAS-CD8::GFP/+ | UAS-ball-RNAi (BL#55330) |
| 2T | y<sup>1</sup> sc* v<sup>1</sup> sev<sup>21</sup>/+ or Y; R71G10-QF2, QUAS-mtdT:3xHA/P{y[+t7.7] v[+t1.8]=TRiP.HMS01723}attP40; Repo-Gal4 UAS-CD8::GFP/+ | UAS-Syx13-RNAi (BL#38525) |
| 2U | y<sup>1</sup> sc* v<sup>1</sup> sev<sup>21</sup>/+ or Y; R71G10-QF2, QUAS-mtdT:3xHA/P{y[+t7.7] v[+t1.8]=TRiP.HMS01756}attP40; Repo-Gal4 UAS-CD8::GFP/+ | UAS-Lac-RNAi (BL#38536) |
| 2V | y<sup>1</sup> v<sup>1</sup>/+ or Y; R71G10-QF2, QUAS-mtdT:3xHA/+; P{y[+t7.7] v[+t1.8]=TRiP.JF01306}attP2/Repo-Gal4 UAS-CD8::GFP | UAS-Art1-RNAi (BL#31348) |
| 2W | y<sup>1</sup> v<sup>1</sup>/+ or Y; R71G10-QF2, QUAS-mtdT:3xHA/+; P{y[+t7.7] v[+t1.8]=TRiP.JF01160}attP2/Repo-Gal4 UAS-CD8::GFP | UAS-Swip-1-RNAi (BL#31585) |
| 2X | y<sup>1</sup> sc* v<sup>1</sup> sev<sup>21</sup>/+ or Y; R71G10-QF2, QUAS-mtdT:3xHA/P{y[+t7.7] v[+t1.8]=TRiP.HMS02429}attP40; Repo-Gal4 UAS-CD8::GFP/+ | UAS-Klc-RNAi (BL#42597) |
| S2A | y<sup>1</sup> v<sup>1</sup>/+ or Y; R71G10-QF2, QUAS-mtdT:3xHA/+; P{y[+t7.7] v[+t1.8]=TRiP.JF01355}attP2/+; OK107-Gal4/+ | UAS-Luciferase-RNAi (BL#31603) |
| S2B | y<sup>1</sup> v<sup>1</sup> /+ or Y; R71G10-QF2, QUAS-mtdT:3xHA/P P{y[+t7.7] v[+t1.8]=TRiP.HMS05812}attP40;; OK107-Gal4/+ | UAS-eRF1-RNAi (BL#67900) |
| 3B | y<sup>1</sup> sc* v<sup>1</sup> sev<sup>21</sup>/+ or Y; R71G10-QF2, QUAS-mtdT:3xHA/ P{y[+t7.7] v[+t1.8]=TRiP.HMC04724}attP40; Repo-Gal4 UAS-CD8::GFP/+ | UAS-CG2712-RNAi (BL#57418) |
| 3C | y<sup>1</sup> v<sup>1</sup>/+ or Y; R71G10-QF2, QUAS-mtdT:3xHA/+; P{y[+t7.7] v[+t1.8]=TRiP.JF01336}attP2/Repo-Gal4, UAS-CD8::GFP | UAS-Mmp1-RNAi (BL#31489) |
| 3D | y<sup>1</sup> v<sup>1</sup>/+ or Y; R71G10-QF2, QUAS-mtdT:3xHA/P{y[+t7.7] v[+t1.8]=TRiP.HMJ23143}attP40; Repo-Gal4, UAS-CD8:GFP/+ | UAS-Mmp2-RNAi (BL#61309) |
| 3F | y<sup>1</sup> v<sup>1</sup>/+ or Y; R71G10-QF2, QUAS-mtdT:3xHA/+; P{y[+t7.7] v[+t1.8]=TRiP.JF01355}attP2/+; OK107-Gal4/+ | UAS-Luciferase-RNAi (BL#31603) |
| 3G | y<sup>1</sup> v<sup>1</sup>/+ or Y; R71G10-QF2, QUAS-mtdT:3xHA/+; P{y[+t7.7] v[+t1.8]=TRiP.JF01336}attP2/+; OK107-Gal4/+ | UAS-Mmp1-RNAi (BL#31489) |
| 3H | y<sup>1</sup> v<sup>1</sup>/+ or Y; R71G10-QF2, QUAS-mtdT:3xHA/P{y[+t7.7] v[+t1.8]=TRiP.HMJ23143}attP40;; OK107-Gal4/+ | UAS-Mmp2-RNAi (BL#61309) |
| 3J | y<sup>1</sup> sc* v<sup>1</sup> sev<sup>21</sup>/+ or Y; R71G10-QF2, QUAS-mtdT:3xHA/ P{y[+t7.7] v[+t1.8]=TRiP.HMC04724}attP40; Repo-Gal4 UAS-CD8::GFP/+ | UAS-CG2712-RNAi (BL#57418) |
| 3K | w/+ or Y; R71G10-QF2, QUAS-mtdT:3xHA/+; {UAS-Mmp1.f1<sup>Δcat</sup>}1/Repo-Gal4, UAS-CD8:GFP | APM 1044 (Andrea Page-McCaw) |
| 3L | +/+ or Y; R71G10-QF2, QUAS-mtdT:3xHA/UAS-Mmp2-RNAi; Repo-Gal4, UAS-CD8:GFP/+ | UAS-Mmp2-RNAi (VDR#107888) |
| S3A | R71G10-QF2, QUAS-mtdT:3xHA/+; Repo-Gal4, UAS-CD8:GFP/+ | Canton S (BL#64349) |
| S3B | y<sup>1</sup> v<sup>1</sup>/+ or Y; R71G10-QF2, QUAS-mtdT:3xHA/P{y[+t7.7] v[+t1.8]=TRiP.HMJ23143}attP40;; OK107-Gal4/+ | UAS-Mmp2-RNAi (BL#61309) |
| S3C | y<sup>1</sup> v<sup>1</sup>/+ or Y; R71G10-QF2, QUAS-mtdT:3xHA/P{y[+t7.7] v[+t1.8]=TRiP.HMJ23143}attP40;; OK107-Gal4/+ | UAS-Mmp2-RNAi (BL#61309) |
| S3D | y<sup>1</sup> v<sup>1</sup>/+ or Y; R71G10-QF2, QUAS-mtdT:3xHA/P{y[+t7.7] v[+t1.8]=TRiP.HMJ23143}attP40;; OK107-Gal4/+ | UAS-Mmp2-RNAi (BL#61309) |
| S3E | y<sup>1</sup> v<sup>1</sup>/+ or Y; R71G10-QF2, QUAS-mtdT:3xHA/P{y[+t7.7] v[+t1.8]=TRiP.HMJ23143}attP40; Repo-Gal4, UAS-CD8:GFP/+ | UAS-Mmp2-RNAi (BL#61309) |
| S4A-B | w/y<sup>1</sup> w<sup>118</sup> or Y; R71G10-QF2, QUAS-mtdT:3xHA/+; 57C10-Gal4/+ | y<sup>1</sup>,w<sup>118</sup> (BL#6598) |
| S4C | w/y<sup>1</sup> w<sup>118</sup> or Y; UAS-mCD4:GFP/+; TIFR-Gal4/+ | TIFR-Gal4 (BL#90392) |
| S4D | w/y<sup>1</sup> w<sup>118</sup> or Y; UAS-mCD4:GFP/+; TIFR-Gal4/P{y[+t7.7] v[+t1.8]=TRiP.JF01336}attP2 | UAS-Mmp1-RNAi (BL#31489) |
| S5B | y,w,hsFlp122, UAS-CD8::GFP/y,w or Y; R71G10-Gal4,G13,Gal80/40A,G13,cn,bw | G13 and 40A are FRTs on 2R and 3R, respectively |
| S5C | y,w,hsFlp122, UAS-CD8::GFP/y,w or Y; R71G10-Gal4,G13,Gal80/40A,G13,mmp2<sup>Δ54-120.C</sup>,cn,bw | G13 and 40A are FRTs on 2R and 3R, respectively |
| S6A | y<sup>1</sup> v<sup>1</sup>/+ or Y; R71G10-QF2, QUAS-mtdT:3xHA/+; P{y[+t7.7] v[+t1.8]=TRiP.JF01355}attP2/+; OK107-Gal4/+ | UAS-Luciferase-RNAi (BL#31603) |
| S6B | y<sup>1</sup> v<sup>1</sup>/+ or Y; R71G10-QF2, QUAS-mtdT:3xHA/+; {UAS-Mmp1.f1<sup>Δcat</sup>}1/+; OK107-Gal4/+ | APM 1044 (Andrea Page-McCaw) |
| S6C | y<sup>1</sup> v<sup>1</sup>/+ or Y; R71G10-QF2, QUAS-mtdT:3xHA/ UAS-Mmp2-RNAi;; OK107-Gal4/+ | UAS-Mmp2-RNAi (VDR#107888) |
| S7A, S9A | y<sup>1</sup> v<sup>1</sup>/+ or Y; R71G10-QF2, QUAS-mtdT:3xHA/+; P{y[+t7.7] v[+t1.8]=TRiP.JF01355}attP2/+; OK107-Gal4/+ | UAS-Luciferase-RNAi (BL#31603) |
| S7B, S9B | y<sup>1</sup> v<sup>1</sup>/+ or Y; R71G10-QF2, QUAS-mtdT:3xHA/+; P{y[+t7.7] v[+t1.8]=TRiP.JF01336}attP2/+; OK107-Gal4/+ | UAS-Mmp1-RNAi (BL#31489) |
| S7C, S9C | y<sup>1</sup> v<sup>1</sup>/+ or Y; R71G10-QF2, QUAS-mtdT:3xHA/P{y[+t7.7] v[+t1.8]=TRiP.HMJ23143}attP40;; OK107-Gal4/+ | UAS-Mmp2-RNAi (BL#61309) |
| S7D, S9D | y<sup>1</sup> v<sup>1</sup>/+ or Y; R71G10-QF2, QUAS-mtdT:3xHA/+; P{y[+t7.7] v[+t1.8]=TRiP.JF01355}attP2/Repo-Gal4, UAS-CD8:GFP | UAS-Luciferase-RNAi (BL#31603) |
| S7E, S9E | y<sup>1</sup> v<sup>1</sup>/+ or Y; R71G10-QF2, QUAS-mtdT:3xHA/+; P{y[+t7.7] v[+t1.8]=TRiP.JF01336}attP2/Repo-Gal4, UAS-CD8:GFP | UAS-Mmp1-RNAi (BL#31489) |
| S7F, S9F | y<sup>1</sup> v<sup>1</sup>/+ or Y; R71G10-QF2, QUAS-mtdT:3xHA/P{y[+t7.7] v[+t1.8]=TRiP.HMJ23143}attP40; Repo-Gal4, UAS-CD8:GFP/+ | UAS-Mmp2-RNAi (BL#61309) |
| 4A | y<sup>1</sup> v<sup>1</sup>/+ or Y; R71G10-QF2, QUAS-mtdT:3xHA/+; P{y[+t7.7] v[+t1.8]=TRiP.JF01336}attP2/Repo-Gal4, UAS-CD8:GFP | UAS-Mmp1-RNAi (BL#31489) |
| 4B | y<sup>1</sup> v<sup>1</sup>/+ or Y; R71G10-QF2, QUAS-mtdT:3xHA/P{y[+t7.7] v[+t1.8]=TRiP.HMJ23143}attP40; Repo-Gal4, UAS-CD8:GFP/+ | UAS-Mmp2-RNAi (BL#61309) |
| 4C | y<sup>1</sup> v<sup>1</sup>/+ or Y; R71G10-QF2, QUAS-mtdT:3xHA/P{y[+t7.7] v[+t1.8]=TRiP.HMJ23143}attP40; P{y[+t7.7] v[+t1.8]=TRiP.JF01336}attP2/Repo-Gal4, UAS-CD8:GFP | UAS-Mmp1-RNAi (BL#31489) + UAS-Mmp2-RNAi (BL#61309) |
| 4E | y<sup>1</sup> v<sup>1</sup> /+ or Y; R71G10-QF2, QUAS-mtdT:3xHA/+; P{y[+t7.7] v[+t1.8]=TRiP.JF01336}attP2/+; OK107-Gal4/+ | UAS-Mmp1-RNAi (BL#31489) |
| 4F | y<sup>1</sup> v<sup>1</sup>/+ or Y; R71G10-QF2, QUAS-mtdT:3xHA/P{y[+t7.7] v[+t1.8]=TRiP.HMJ23143}attP40;; OK107-Gal4/+ | UAS-Mmp2-RNAi (BL#61309) |
| 4G | y<sup>1</sup> v<sup>1</sup>/+ or Y; R71G10-QF2, QUAS-mtdT:3xHA/P{y[+t7.7] v[+t1.8]=TRiP.HMJ23143}attP40; P{y[+t7.7] v[+t1.8]=TRiP.JF01336}attP2/+; OK107-Gal4/+ | UAS-Mmp1-RNAi (BL#31489) + UAS-Mmp2-RNAi (BL#61309) |
| 4I | y<sup>1</sup> v<sup>1</sup>/+ or Y; R71G10-QF2, QUAS-mtdT:3xHA/+; P{y[+t7.7] v[+t1.8]=TRiP.JF01336}attP2/Repo-Gal4, UAS-CD8:GFP; OK107-Gal4/+ | UAS-Mmp1-RNAi (BL#31489) |
| 4J | y<sup>1</sup> v<sup>1</sup>/+ or Y; R71G10-QF2, QUAS-mtdT:3xHA/P{y[+t7.7] v[+t1.8]=TRiP.HMJ23143}attP40; Repo-Gal4 UAS-CD8:GFP/+; OK107-Gal4/+ | UAS-Mmp2-RNAi (BL#61309) |
| 4K | y<sup>1</sup> v<sup>1</sup>/+ or Y; R71G10-QF2, QUAS-mtdT:3xHA/P{y[+t7.7] v[+t1.8]=TRiP.HMJ23143}attP40; P{y[+t7.7] v[+t1.8]=TRiP.JF01336}attP2/Repo-Gal4, UAS-CD8:GFP; OK107-Gal4/+ | UAS-Mmp1-RNAi (BL#31489) + UAS-Mmp2-RNAi (BL#61309) |
| S8A | y<sup>1</sup> w<sup>118</sup> /+ or Y; R71G10-QF2, QUAS-mtdT:3xHA/+; +/Alrm-Gal4 UAS-CD8::GFP | y<sup>1</sup> ,w<sup>118</sup> (BL#6598) |
| S8B | y<sup>1</sup> v<sup>1</sup>/+ or Y; R71G10-QF2, QUAS-mtdT:3xHA/+; P{y[+t7.7] v[+t1.8]=TRiP.JF01336}attP2/Alrm-Gal4, UAS-CD8:GFP | UAS-Mmp1-RNAi (BL#31489) |
| S8C | y<sup>1</sup> v<sup>1</sup>/+ or Y; R71G10-QF2, QUAS-mtdT:3xHA/P{y[+t7.7] v[+t1.8]=TRiP.HMJ23143}attP40; Alrm-Gal4, UAS-CD8:GFP/+ | UAS-Mmp2-RNAi (BL#61309) |

### Drosophila MMPs are required in KCs and glia for γ axon pruning

Many of the ‘positive hits’ in the screen are promising directions for further exploration. Interestingly, two genes - Syntaxin13 (Syx13) and Matrix Metalloproteinase 2 (Mmp2) - emerged as required for pruning in both neurons and glia. We decided to delve deeper into Mmp2, as a previous, CRISPR-based screen done by our lab also found it to be required in γ-KCs for axon pruning (Meltzer et al., 2019).

MMPs are extracellular proteases that cleave extracellular matrix (ECM) components (Page-McCaw et al., 2007; Sreesada et al., 2025). In the mouse there are 23 MMPs, while the zebrafish genome encodes 25 MMPs. The *Drosophila* genome, in contrast, encodes only two MMPs – Mmp1 and Mmp2 – which together have 13 predicted isoforms, most of which are likely secreted, while each Mmp also has a glycosylphosphatidylinositol (GPI)-anchored isoform (LaFever et al., 2017). The significantly reduced complexity makes *Drosophila* MMPs an ideal model system to uncover novel insights into their roles in neuronal remodeling. Our transcriptional datasets indicate that both MMPs are dynamically expressed in γ-KCs, as well as in larval astrocytes (Fig. 3A). We thus decided to KD Mmp1 in both KCs and glia, and compare it to Mmp2 (Fig. 3B-I). Blinded ranking by two independent investigators (Fig. S3) established that the pruning defect induced by KD of Mmp1 or Mmp2 is significant compared to controls in glia (Fig. 3B-E). KD of Mmp1 in KCs results in a mild pruning defect, not reaching statistical significance, while knockdown of Mmp2 is significant compared to controls (Fig. 3F-I). Importantly, we validated the efficiency of the Mmp1 RNAi line in reducing protein levels using an Mmp1 antibody (Fig. S4). The specific Mmp2 RNAi line that we used was previously demonstrated to efficiently reduce Mmp2 protein levels (Harmansa et al., 2023).

**Figure 3.**
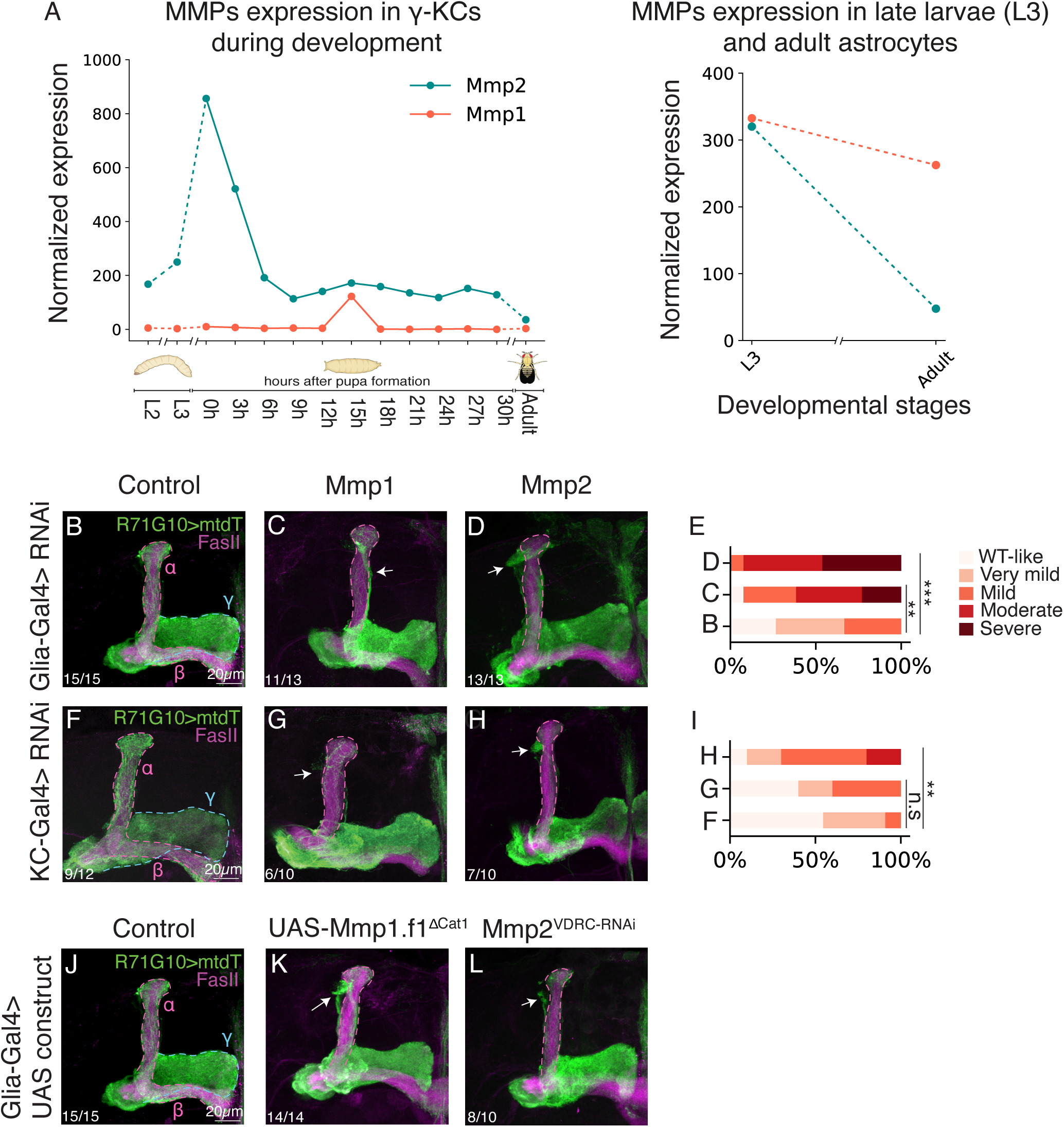
*Drosophila* MMPs are required in KCs and glia for γ-axon pruning. (A) Normalized gene expression of Mmp1 (orange) and Mmp2 (cyan) in developing γ-KCs (left; based on Alyagor et al., 2018) and in larval/adult astrocytes (right; based on Marmor-Kollet et al., 2023). (B-D, F-H) Confocal z-projections of adult MBs in which γ-KCs are labeled by R71G10-QF2-driven QUAS-mtdT (green). RNAi species targeting control genes (B, F), Mmp1 (C, G) or Mmp2 (D, H) are expressed in all glia using Repo-Gal4 (B-D), or in all KCs using OK107-Gal4 (F-H). Control in B is RNAi targeting CG2712 (see Fig. 1D), and in F is RNAi targeting Luciferase. (E, I) Quantification of the pruning defect phenotypes, ranging from 1 (WT-like phenotype) to 5 (severe pruning defect). Genotypes are indicated with the letter of the corresponding image within this figure. B vs. C p=0.001, B vs. D p=0.0001, F vs. G p=0.289, F vs. H p=0.009. *** = p<0.001, * = p<0.05, n.s = not significant. For examples of quantification scores as well as ranking comparison see Fig. S3. (J-L) Confocal z-projections of adult MBs in which γ-KCs are labeled by R71G10-QF2-driven QUAS-mtdT (green). Indicated UAS transgenes are driven by Repo-Gal4 Control in J is RNAi targeting CG2712 (see Fig. 1D). Arrows indicate unpruned axons. The γ lobe is outlines ib blue and the α/β lobes in magenta. FasII antibody (magenta) strongly labels α/β-axons and weakly labels γ-axons. Scale bar corresponds to 20µm.

We next used CRISPR/Cas9 to generate a novel, predicted null *mmp2* mutant allele. Due to its (expected) lethality, we employed MARCM (Lee & Luo, 1999) to generate homozygous mutant γ-KC clones. However, *mmp2*^Δ54-120.PC^ clones did not present pruning defects (Fig. S5), consistent with its predicted function as a predominantly secreted protein, and therefore a non-cell autonomous role. Thus, to further validate our findings, we tested a second RNAi line targeting Mmp2, which similarly showed pruning defects when driven in glia (Fig. 3L), and a very mild defect in KCs (Fig. S6C).

To further explore the requirement of Mmp1, we tested a dominant negative (DN) variant previously generated by the Page-McCaw lab (Glasheen et al., 2009). When expressed in glia, Mmp1-DN resulted in a severe γ-axon pruning defect (Fig. 3K), while expressing it in KCs results in a mild pruning defect (Fig. S6B), reinforcing the reduced requirement of Mmp1 in γ-KCs compared to glia.

Notably, the initial growth of γ-KC axons, as evident at the onset of pupariation, is unaffected by KD of either Mmp1 or Mmp2, in neither KCs nor glia (Fig. S7), highlighting their specific role in the axon remodeling phase.

Taken together, our data suggests that while both MMPs are required in glia and in KCs for γ-axon remodeling.

### Analysis of combinatorial MMP knockdown in both KCs and glia

Since most Mmp1 and Mmp2 isoforms are secreted, we speculated that both are secreted in parallel from KCs and glia to promote axon pruning. Therefore, we preformed different single and double KD combinations of Mmp1 and/or Mmp2 in KCs and/or glia (Fig. 4). Our data and quantification (by two independent investigators, see Methods and Fig. S3) draw two main conclusions: first, while both Mmp1 and Mmp2 are required for pruning, using the available drivers and RNAi reagents, the observed phenotypes of Mmp2 KD are more severe in all conditions. Second, while Mmp1/2 expression from both KCs and glia is required for pruning of γ-axons, the main source is likely glia, as KD of a single or double MMP in both glia and neurons did not further exacerbate the pruning defect severity compared to the respective KD in glia only (Fig. 4, quantified in M-O).

**Figure 4.**
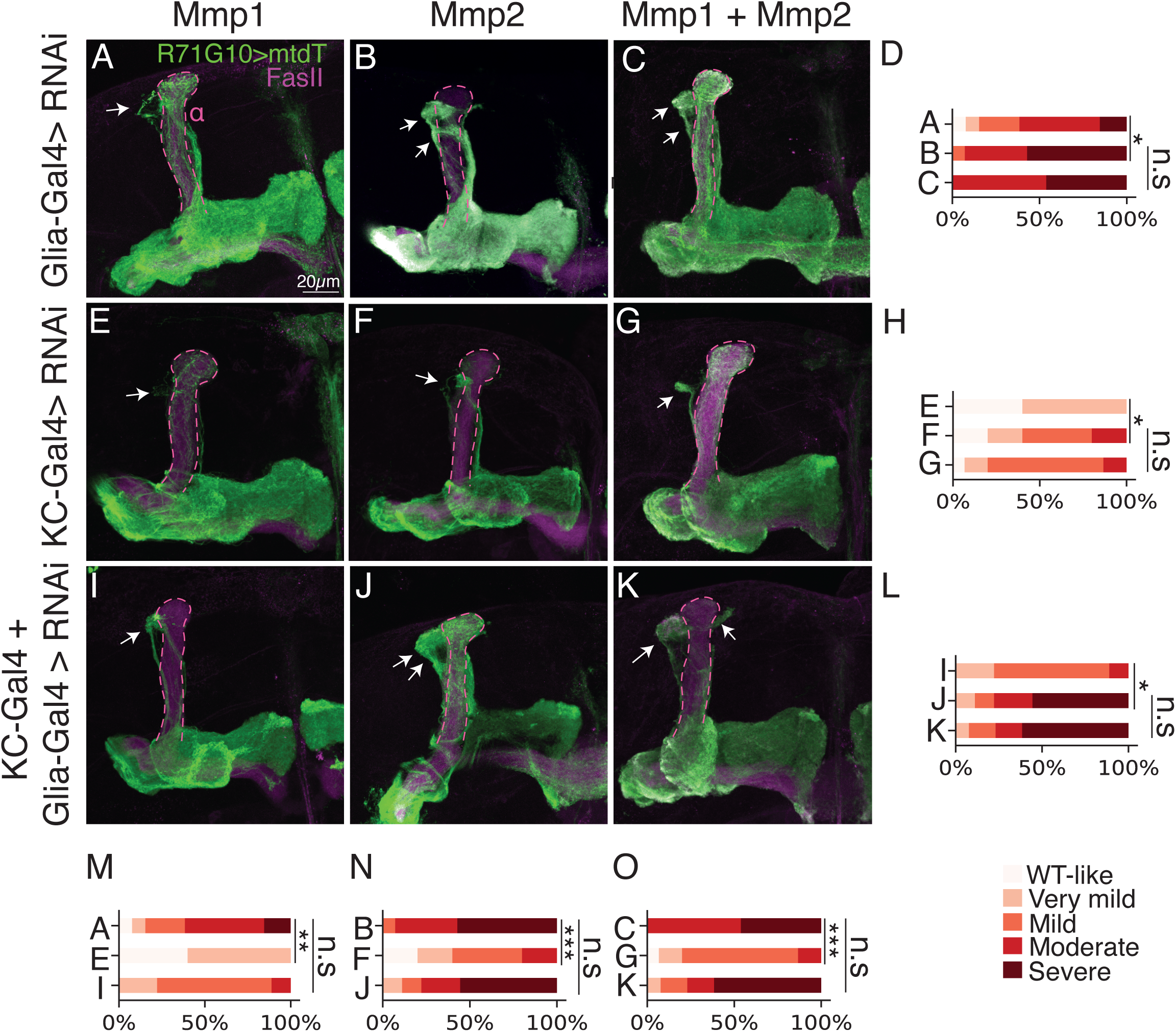
**Analysis of combined MMP knockdown in both KCs and glia.** (A-C, E-G, I-K) Confocal z-projections of adult MBs in which γ-KCs are labeled by R71G10-QF2- driven QUAS-mtdT (green), and UAS-RNAi species targeting Mmp1 (A, E, I), Mmp2 (B, F J), or both (C, G, K) are driven by Repo-Gal4 (A-C), OK107-Gal4 (E-G), or both simultaneously (I-K). FasII antibody (magenta) strongly labels α/β-axons and weakly labels γ-axons. Arrows indicate unpruned axons. The α-lobe is outlined in magenta. (D, H, L-O) Quantification of the pruning defect phenotypes, ranging from 1 (WT-like phenotype) to 5 (severe pruning defect). Genotypes are indicated with the letter of the corresponding image within this figure. For quantification score examples and ranking comparison see Fig. S3. *** = p<0.001, * = p<0.05, n.s = not significant. A vs. B p=0.023, B vs. C p=0.860, E vs. F p=0.046, F vs. G p=0.719, I vs. J p=0.023, J vs. K p=0.943, A vs. E p=0.001, A vs. I p=0.092, B vs. F p=0.0009, B vs. J p=0.860, C vs. G p=0.0003, C vs. K p=0.943.

Due to the major glial contribution, combine with the known astrocytic involvement in remodeling and the expression data (Fig. 3A), we decided to also test KD using the astrocyte-specific driver alrm-Gal4, which indeed resulted in mild pruning defect in the case of Mmp2 (Fig. S8). The mild phenotype can be explained by weaker Gal4 expression compared to Repo-Gal4, or due to involvement of additional glial subtypes. Unfortunately, our attempts to analyze other glial subtypes were abruptly terminated (see acknowledgements).

Taken together, our results suggest that while both MMPs affect pruning likely via secretion from both neurons and glia, the strongest requirement for MB γ axon pruning comes from MMPs secretion by glia - at least partially by astrocytes.

## Discussion

Four decades ago, Feinberg hypothesized that SCZ is caused by aberrant synaptic pruning during adolescence: “too many, too few or the wrong synapses are eliminated” (Feinberg, 1982). In recent years, accumulating findings are providing experimental support of this hypothesis (Keshavan et al., 2020; Sekar et al., 2016; Sellgren et al., 2019). In this work, we aimed to further explore the molecular association of SCZ-related genes with defective neuronal remodeling, by utilizing the stereotypic remodeling of the *Drosophila* MB.

In a genetic LOF screen for *Drosophila* homologs of human genes that contain SNPs associated with SCZ, about 40% of the genes we screened within the neurons (11 out of 29) were found to have a role in γ-axon pruning. Additionally, 7 out of 8 genes we screened within the glia were found to have a role in γ-axon pruning. This high rate of ‘positive hits’ strengthens the link between SCZ-associated genes and defects in developmental neuronal remodeling and emphasizes glia’s major role. Notably, we focused on genes that are upregulated prior to pruning, but future efforts to overexpress genes that are downregulated prior to pruning might also reveal interesting insights.

While many of the ‘positive hits’ are fascinating candidates for further research, in this study we decided to focus on the role of MMPs in neuronal remodeling, for several reasons. First, Mmp2 was found to be required in both glia and neurons for γ-axon pruning. This is of particular interest due to the speculated role of glia in abnormal synaptic pruning in SCZ patients. Second, previous reports have highlighted the link between MMPs and the pathophysiology of SCZ. Specifically, elevated levels of MMPs, most prominently Mmp9, were shown to correlate with SCZ risk and cognitive impairment

(Dickerson et al., 2023; Kudo et al., 2020; Schoretsanitis et al., 2021; Seitz-Holland et al., 2022). While it has been suggested that this correlation is via MMP-dependent changes in dendritic spine morphology (Lepeta et al., 2017), the precise mechanism remains poorly understood. Of note, the *Drosophila* MMPs are evolutionary conserved (LaFever et al., 2017; Llano et al., 2002), highlighting their potential relevance and providing a simplified and powerful model system for mechanistic exploration.

MMPs are known for their roles in tissue remodeling by cleaving substrates in the ECM (Page-McCaw et al., 2007), and are well studied in the cancer field (Cox, 2021; Overall & López-Otín, 2002), but their mechanistic function in the developing nervous system is less characterized (e.g., Broadie et al., 2011; Dear et al., 2016; Gore et al., 2021). MMPs are known to be secreted by neurons and glia, and are thought to facilitate synaptic remodeling of dendritic spines (Huntley, 2012). In adult rodent brains, pharmacologically- (Szklarczyk et al., 2002) or injury- (Pijet et al., 2019) induced dendritic remodeling is suggested to occur via Mmp9. In the developmental remodeling of *Drosophila* dendritic arborization (da) neurons, both Mmp1 and Mmp2 were shown to be required for the elimination of severed dendritic branches in a non-cell autonomous manner (Kuo et al., 2005). Their source, as well as the mechanism by which they promote dendrite removal, remain unknown (the authors speculate they are secreted from phagocytic cells – at the time they suggested blood cells, but given more recent studies from the same lab, epidermal cells are more likely; Han et al., 2014). Interestingly, MMPs were also shown to mediate dendrite reshaping of da neurons in the mature fly via local degeneration of the basement membrane, further reinforcing their significance in refining the mature nervous system (Yasunaga et al., 2010). Finally, it was shown that Mmp1 is upregulated in astrocytes after traumatic brain injury (Li et al., 2024), and in ensheathing glia following ventral nerve cord injury to promote axonal debris clearance (Purice et al., 2017).

The mechanism by which MMPs promote remodeling of MB γ-axons, and specifically their substrates in this context, remain to be uncovered. Since the main known role of MMPs is cleaving ECM substrates, they may act as ‘path cleaners’ for the migration of cells and molecules (Page-McCaw et al., 2007). This function can be relevant to axon pruning in several, non-mutually exclusive ways. One option is that MMPs allow the migration of Myoglianin (Myo), a *Drosophila* TGF-β, in the ECM. Myo is part of the pruning initiation cascade and was shown to be secreted by cortex glia and astrocytes (Awasaki et al., 2011) to eventually promote expression of ecdysone receptor (EcR)-B1, a key regulator of remodeling (Yu et al., 2013; Zheng et al., 2003). Moreover, it was shown that the human Mmp9 can proteolytically activate latent TGF-β (Yu & Stamenkovic, 2000). However, we found that the expression of EcR-B1 in γ-KCs in unaffected by glia or neuronal KD of Mmp1/Mmp2 (Fig. S9). Another option is that MMPs facilitate astrocyte infiltration into the γ-axon bundle at the onset of pruning, that was shown to be required for defasciculation (Marmor-Kollet et al., 2023), and subsequent engulfment of the axonal debris (Hakim et al., 2014; Tasdemir-Yilmaz & Freeman, 2014). Unfortunately, testing this hypothesis requires three binary systems and is beyond the scope of this study. MMPs were also shown to degrade adhesion molecules (Page-McCaw et al., 2007), and our lab previously showed that the adhesion molecule Fasciclin II (FasII) must be downregulated for pruning to occur properly (Bornstein et al., 2015). It is thus possible that MMPs cleave FasII, or other adhesion molecules, to promote axon pruning. Since most *Drosophila* MMP isoforms are secreted (LaFever et al., 2017), together with the finding that they are required from both neuronal and glial origin, it is possible that MMPs are also secreted from additional cells within or around the MB circuit. All the above-mentioned options are promising directions for future research, in hopes of uncovering the mechanisms by which MMPs promote neuronal remodeling.

Notably, the main hypothesis in the literature is that SCZ is associated with increased synapse pruning, while in our study KD of SCZ-associated genes resulted in inhibition of pruning, which may seem counter-intuitive at first. However, SNPs can result in loss- but also gain-of-function, which could account for this apparent conflict. In fact, it was previously shown that the SCZ-associated SNP in Mmp9 is in the 3’ UTR, and affects Mmp9 mRNA folding which eventually results in higher Mmp9 compared to WT (Lepeta et al., 2017).

We recognize that the current study explores pruning of long stretches of axons, while SCZ is mainly associated with defective synapse pruning. More generally, flies could never model the complexities associated with human psychiatric conditions, nor directly offer therapeutic targets. Nonetheless, this simple and genetically accessible model enables us to uncover conserved neurodevelopmental principles and concepts.

Understating the neurodevelopmental role of MMPs may provide insights into brain ECM dynamics throughout development, and specifically during neuronal remodeling.

## Materials and methods

### Gene selection for the screen

Human genes that contain SNPs associated with SCZ were collected from two studies: PGC2 & CLOZUK (Wu et al., 2017; Supplemental excel file 1). These genes were converted to their *Drosophila melanogaster* homologs using HumanMine (Supplemental excel file 1). In cases of multiple fly homologs for a single human gene, we decided to include all of them.

### Drosophila melanogaster rearing and strains

All fly strains were reared under standard laboratory conditions at 25°C on molasses-containing food. Males and females were chosen at random. Unless specifically stated otherwise, the relevant developmental stage is adult, which refers to 3-5 days post eclosion. The RNA lines used for the screen are detailed in Table 1. The following lines were obtained for the Bloomington Drosophila Stock Center (BDSC): Luciferase RNAi (used as control for RNAi experiments; #31603), CG2712 RNAi (used as control for RNAi experiments; #57418), Mmp1 RNAi (#31489), Alrm-Gal4 (#67032), Repo-Gal4 (#7415), OK107-Gal4 (#854), TIFR-Gal4 (#90392), QUAS-mtdTomato-3xHA (#30004), Mmp2-gRNA (#82521). R71G10-QF2 was previously generated by our lab (Bornstein et al., 2021). R71G10-Gal4 on the 2^nd^ chromosome was previously generated by our lab (Alyagor et al., 2018)A second Mmp2 RNAi line was obtained from the Vienna Drosophila Resource Center (VDRC; #107888). Line UAS-Mmp1.f1^ΔCat1^ was a generous gift from Andrea Page-McCaw. See full genotypes (ordered by specific figure panels) in Table 2.

### Immunohistochemistry and imaging

*Drosophila* brains were dissected in cold ringer solution and placed on ice. Followed by fixation in 4% paraformaldehyde (PFA) solution for 20 min in room temperature (RT). Fixed brains were then washed with PB supplemented with 0.3% Triton X-(PBT) – 3 times, 20 min each. Next the brains were blocked for non-specific staining using 5% heat inactivated normal goat serum (NGS). Antibody staining was performed as follows: primary antibodies (4°C, overnight), PBT washes (X3 in RT, 20 min each), secondary antibodies (RT, 2h), PBT washes (X3 in RT, 20 min each), mounting in SlowFade (Invitrogen). Primary antibodies included mouse anti-FasII 1:25 (1D4; Developmental Studies Hybridoma Bank; DSHB), mouse anti-EcRB1 1:25 (AD4.4; DSHB), rat anti-HA 1:250 (Sigma Aldrich, 11867423001) and mouse anti-Mmp1 1:20 (mix of: 3A6B4, 3B8D12 and 5H7B11; DSHB). Secondary antibodies included Alexa fluor 647 goat anti-mouse 1:300 (A-21236; Invitrogen) and Alexa fluor 568 goat anti-rat 1:300 (A-21247; Invitrogen). All brains were imaged on a Zeiss LSM980 confocal microscope. The images were processed using ImageJ 2.14.0 (NIH).

### Generation of the mmp2 mutant allele

The *mmp2* mutant allele named Δ54-120.C was generated by crossing a gRNA-line, which harbors two distinct gRNA sequences both targeting the 5’ CDS region common to all 3 Mmp2 isoforms (cloned into pCFD4; BDSC #82521, WKO.3-C7), to a nanos-Cas9- expressing fly. The recovered indel is a deletion of 67bp (54-120 in the CDS of isoform C) – predicted to induce a premature stop codon (after 77 out of 606 amino acids in isoform C). Unfortunately, this mutant (alongside many others) was lost on June 15^th^ (see acknowledgements).

The following primers were used to sequence the indel mutation: F: GCATTCAATGCTGCCACAAA R: CATTTCATCATCGACGTCGT

### Generation of MARCM clones

γ-KC neuroblast clones were generated using the MARCM strategy (Lee & Luo, 1999). Newly hatched larvae (approximately 24h after egg laying) were heat-shocked for 1 hour in 37°C. Adult brains were dissected for further analysis.

### Quantification and statistical analysis

Blind ranking of the pruning phenotypes for Fig. 3 and Fig. 4 were performed, for all experiments, by two independent investigators who reviewed the Z-projections. Ranking scores: 1 = WT-like phenotype, 2 = very mild pruning defect (PD), 3 = mild PD, 4 = moderate PD, 5 = severe PD (Fig. S3A-E). The two sets of ranking were compared using Wilcoxon signed rank test and found to be statistically insignificant (Fig. S3F-G).

A non-parametric statistical analysis was preformed to analyze the ranking (Fig. 3 and Fig. 4): Kruskal Wallis H test followed by a post hoc Mann Whitney U test with FDR correction. *** = p value<0.001, ** = p value<0.01, * = p value<0.05, N.S = not significant. Specific p values are indicated in the figure legends.

## Acknowledgements

We thank Andrea Page-McCaw (Vanderbilt University) for kindly sharing the Mmp1 dominant negative strain, and the Bloomington Drosophila Stock Center for reagents. Monoclonal antibodies were obtained from the Developmental Studies Hybridoma Bank developed under the auspices of the NICHD and maintained by the University of Iowa. We thank R. Rothkopf for assistance with statistics. This work was supported by European Research Council (ERC) advanced grant #<u>101054886</u> “NeuRemodelBehavior,” and by the ISF-NSFC joint research program (grant no. <u>2573/18</u>). O.S is an incumbent of the Prof. Erwin Netter Professorial Chair of Cell Biology. On the night of June 15^th^ 2025, our lab was directly hit by an Iranian missile and was completely destroyed. Unfortunately, this affects our abilily to provide reagents and limited the scope of our revisions.

## Author contributions

S.K. conceptualized the project, designed, performed and analyzed experiments, performed bioinformatic and statistical analyses, and wrote the manuscript. H.M. designed and performed experiments, interpreted results, and wrote the manuscript. N.M-K. designed experiments and assisted with bioinformatic analyses. O.S. led the project, designed experiments, interpreted results, wrote the manuscript, and procured funding.

