## Supplementary material for "A *Drosophila* screen of schizophrenia-related genes highlights the requirement of neural and glial matrix metalloproteinases for neuronal remodeling": Fig. S1

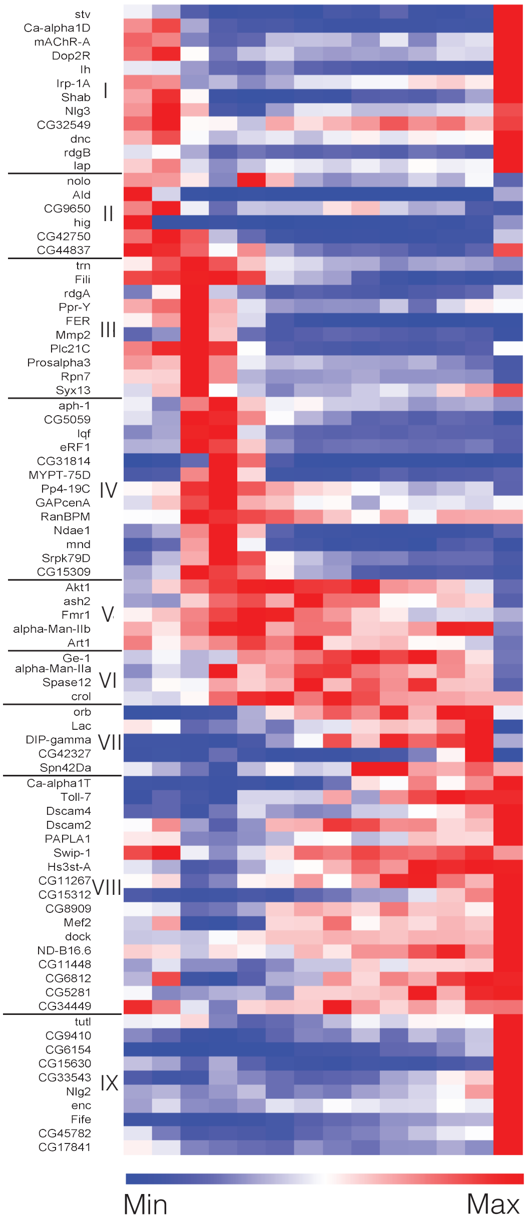
stv
Ca-alpha1D
mAChR-A
Dop2R
Ih
Irp-1A
Shab
Nlg3
CG32549
dnc
rdgB
lap
I
nolo
Ald
CG9650
hig
CG42750
CG44837
II
trn
Fili
rdgA
Ppr-Y
FER
Mmp2
Plc21C
Prosalpha3
Rpn7
Syx13
III
aph-1
CG5059
lqf
eRF1
CG31814
MYPT-75D
Pp4-19C
GAPcenA
RanBPM
Ndae1
mnd
Srpk79D
CG15309
IV
Akt1
ash2
Fmr1
alpha-Man-IIb
Art1
V
Ge-1
alpha-Man-IIa
Spase12
crol
VI
orb
Lac
DIP-gamma
CG42327
Spn42Da
VII
Ca-alpha1T
Toll-7
Dscam4
Dscam2
PAPLA1
Swip-1
Hs3st-A
CG11267
CG15312
CG8909
Mef2
dock
ND-B16.6
CG11448
CG6812
CG5281
CG34449
VIII
tutl
CG9410
CG6154
CG15630
CG33543
Nlg2
enc
Fife
CG45782
CG17841
IX
Min
Max

**Figure S1. *Drosophila* homologs of SCZ-related genes are dynamically expressed in developing γ-KCs**

A full heatmap (including gene symbols) representing the expression of 82 dynamically expressed genes along γ-KC development, clusters I-IX (based on Alyagor et al., 2018; Supplemental excel file 1). Each row represents a gene, with red and blue indicating high and low relative expression, respectively. The genes in clusters III-VI were selected as candidates for screening (Fig. 1C; see Table 1 for the list of *Drosophila* lines).
