## Supplementary material for "A *Drosophila* screen of schizophrenia-related genes highlights the requirement of neural and glial matrix metalloproteinases for neuronal remodeling": Fig. S2

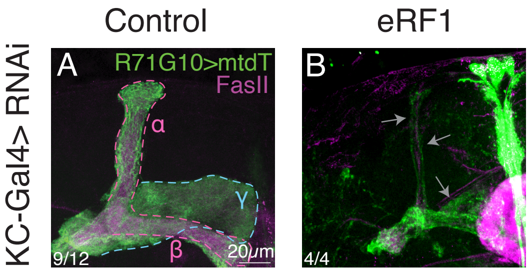

Control
eRF1
KC-Gal4> RNAi
A
R71G10>mtdT
FasII
α
γ
β
20μm
9/12
B
4/4

**Figure S2. KD of a *Drosophila* homolog of eRF1 causes abnormal mushroom body morphology**

(A-B) Confocal z-projections of adult MBs, either control RNAi targeting luciferase (A) or RNAi targeting eukaryotic translation release factor 1 (eRF1; B), expressed OK107-Gal4. R71G10-QF2 drives expression of membranal tandem tomato (mtdT; green) in γ-KCs and stochastically in α/β-KCs. FasII antibody (magenta) strongly labels α/β-KCs and weakly labels γ-KCs. Gray arrows indicate abnormal structures. Scale bar corresponds to 20µm. The number of MBs (each from an individual brain) showing the presented phenotype out of the total n for each genotype is indicated.
