## Supplementary material for "A *Drosophila* screen of schizophrenia-related genes highlights the requirement of neural and glial matrix metalloproteinases for neuronal remodeling": Fig. S3

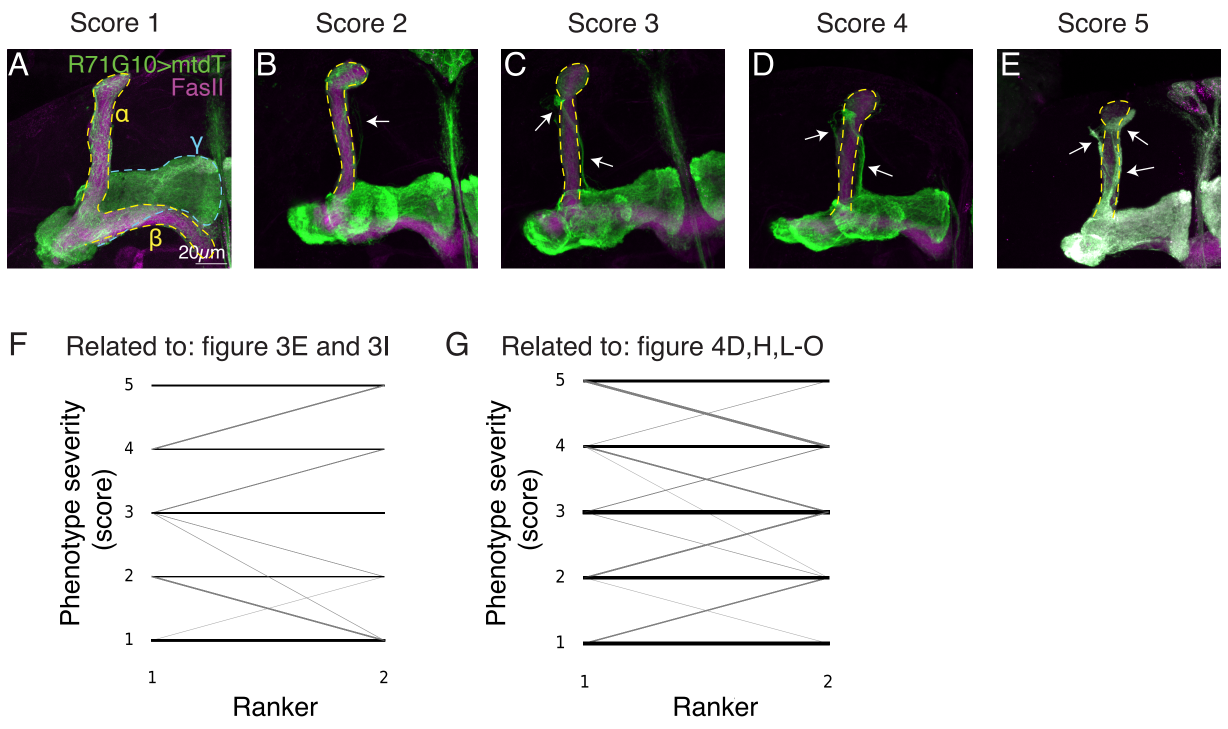

Score 1
Score 2
Score 3
Score 4
Score 5
A
R71G10>mtdT
FasII
α
γ
β
20μm
B
C
D
E
F
Related to: figure 3E and 3I
Phenotype severity (score)
Ranker
G
Related to: figure 4D,H,L-O
Phenotype severity (score)
Ranker

**Figure S3. Pruning defect scores and comparison of ranking**

(A-E) Confocal z-projections of adult MBs demonstrating the various degrees of pruning defects (for full list see Table 2). Arrows indicate unpruned axons. Scale bar corresponds to 20μm.

(F,G) Spaghetti plots comparing the scores for each image as ranked by two independent investigators. Each line represepnts a single image. The thicker the line the more frequent the event. (F) is related to the ranking in Figure 3E,I (Wilcoxon signed-rank test W=162.5, p=0.714); (G) is related to the ranking in Figure 4D,H,L-O (Wilcoxon signed-rank test W=483, p=0.475).
