## Supplementary material for "A *Drosophila* screen of schizophrenia-related genes highlights the requirement of neural and glial matrix metalloproteinases for neuronal remodeling": Fig. S4

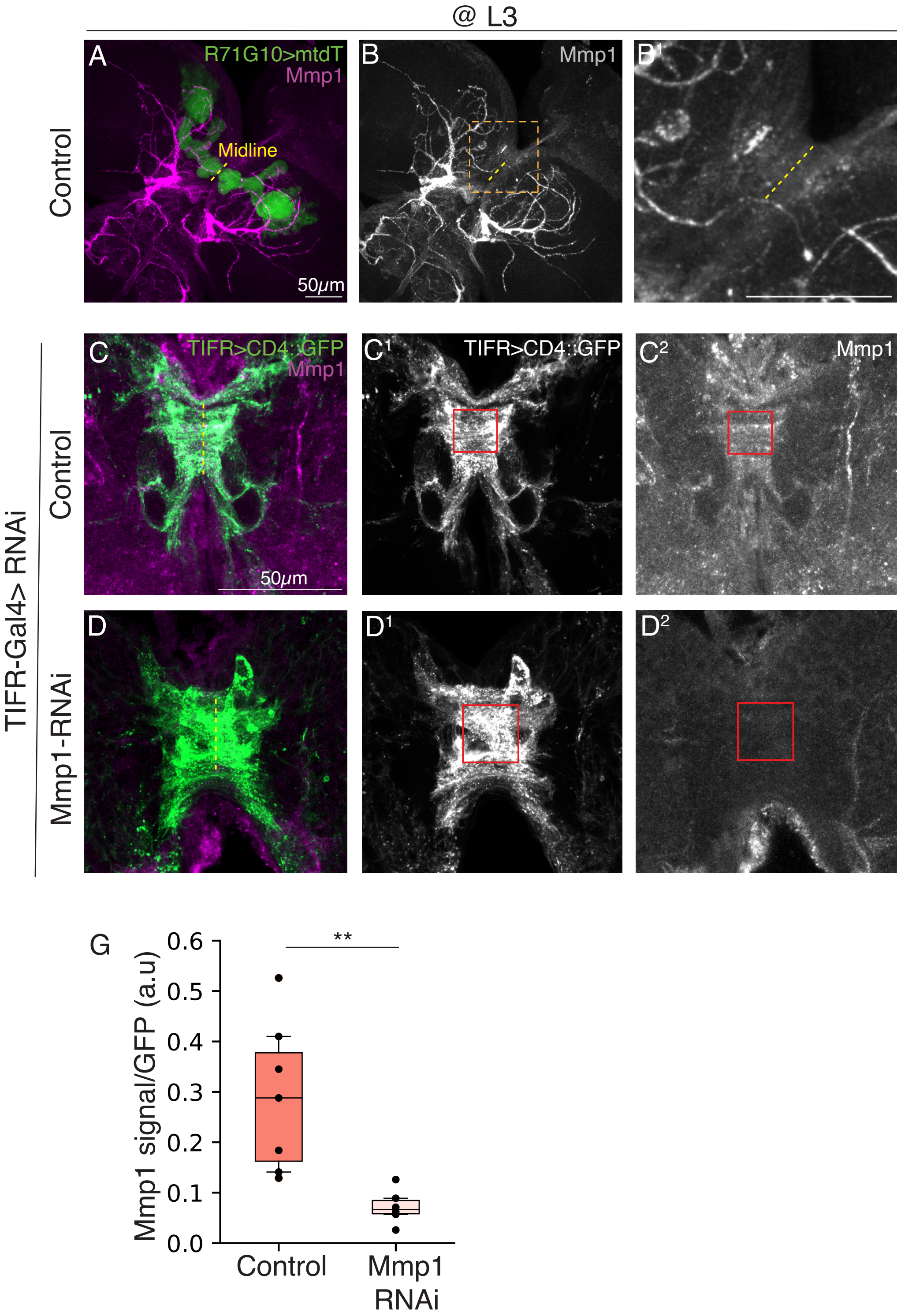

@ L3
A
R71G10>mtdT
Mmp1
Midline
50μm
Control
B
Mmp1
B1
C
TIFR>CD4::GFP
Mmp1
50μm
C1
TIFR>CD4::GFP
C2
Mmp1
D
D1
D2
TIFR-Gal4> RNAi
Control
Mmp1-RNAi
G
Mmp1 signal/GFP (a.u)
0.0
0.1
0.2
0.3
0.4
0.5
0.6
**
Control
Mmp1 RNAi

**Figure S4. Mmp1 RNAi reduces Mmp1 protein levels**

(A-B) Confocal z-projections of L3 MBs in which R71G10-QF2-drives the expression of membranal tandem tomato (mtdT; green) in γ-KCs, and Mmp1 is labeled using an antibody mix (A; magenta, B; gray). The brain midline is marked in a yellow dashed line. We were unable to identify the strongly stained structures, however, as can be seen in the magnified view (the area in the orange box in B) in B1, there is increased Mmp1 staining in the midline region, which potentially correlates with the transient interhemispheric fibrous ring (TIFR) structure. Thus, we decided to test the RNAi using a TIFR-specific driver (see C-D below). Scale bar corresponds to 50µm (A-B).

(C-D) Confocal sub-z-projections of L3 MBs in which TIFR-Gal4 is driving mCD4::GFP (green/grayscale). Mmp1 (Magenta/grayscale) is labeled using an antibody mix. Control (y,w; C) or Mmp1-RNAi (D) is expressed in using TIFR-Gal4. Scale bar corresponds to 50µm. Red outlined boxes define the ROI used to extract mean fluorescence intensity.

(E) Quantification of Mmp1 antibody mean fluorescence intensity within the ROI (red boxes in C-D) divided by GFP mean fluorescence intensity. Two tailed Student's t-test, $p=0.002$. Box plots show the median (horizontal line), interquartile range (box), 10th and 90th percentiles (whiskers), and all individual data points (black dots).
