## Supplementary material for "A *Drosophila* screen of schizophrenia-related genes highlights the requirement of neural and glial matrix metalloproteinases for neuronal remodeling": Fig. S5

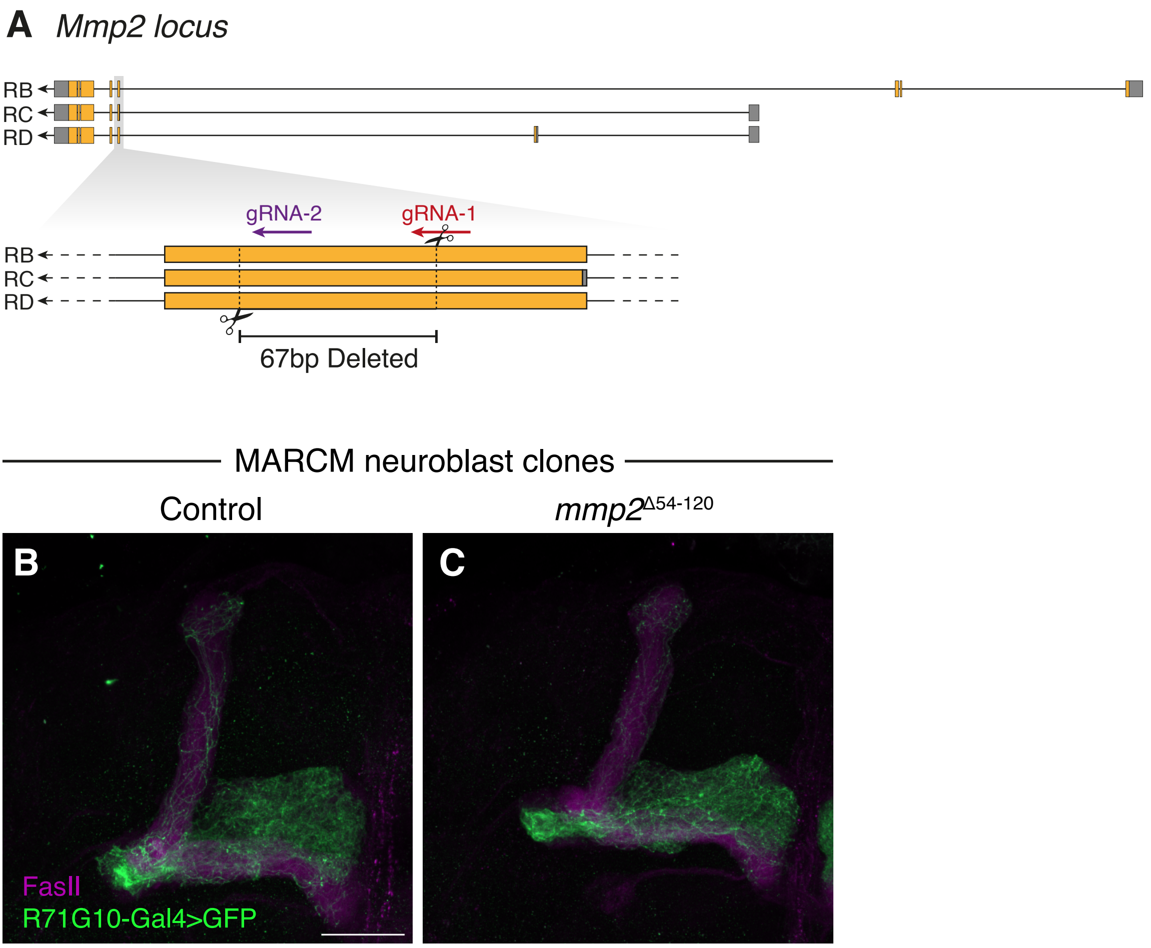

A
Mmp2 locus
RB
RC
RD
gRNA-2
gRNA-1
RB
RC
RD
67bp Deleted
MARCM neuroblast clones
Control
mmp2Δ54-120
B
C
FasII
R71G10-Gal4>GFP

**Figure S5. Clonal analyses of a novel Mmp2 null allele**

(A) Scheme depicting the genomic locus of Mmp2, the location of gRNA sequences and the deleted segment of 67bp (correlating to 54-120 bp of isoform C). Gray and orange represent non-coding and coding exons, respectively; Lines represent introns.

(B-C) Confocal z-projections of γ-KC MARCM neuroblast clones in adult MBs, either control (B) or homozygous mutant for the *mmp2*$^{\Delta 54\text{-}120.C}$ allele (C). Clones are labelled by membranal GFP (CD8::GFP; green) driven by the γ-specific driver R71G10-Gal4. Magenta is FasII staining, which strongly labels α/β-axons and weakly labels γ-axons. Scale bar is 30μm.
