## Supplementary material for "A *Drosophila* screen of schizophrenia-related genes highlights the requirement of neural and glial matrix metalloproteinases for neuronal remodeling": Fig. S6

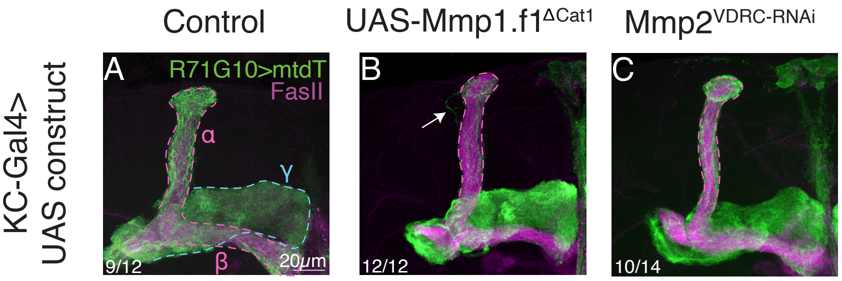

Control
UAS-Mmp1.f1ΔCat1
Mmp2VDRC-RNAi
KC-Gal4>
UAS construct
A
R71G10>mtdT
FasII
α
γ
β
9/12
20μm
B
12/12
C
10/14

**Figure S6. Additional Mmp1 and Mmp2 phenotypes**

(A-C) Confocal z-projections of adult MBs in which γ-KCs are labeled by R71G10-QF2-driven QUAS-mtdT (green), and control (Luciferase-RNAi; A), Mmp1-DN (Mmp1.f1$^{DCat1}$; B) or Mmp2-RNAi (VDRC #107888; C) are expressed in all KCs using OK107-Gal4.

Arrows indicate unpruned axons. The γ lobe is outlines in blue and the α/β lobes are outlined in magenta. FasII antibody (magenta) strongly labels α/β-axons and weakly labels γ-axons. Scale bar corresponds to 20μm.
