## Supplementary material for "A *Drosophila* screen of schizophrenia-related genes highlights the requirement of neural and glial matrix metalloproteinases for neuronal remodeling": Fig. S7

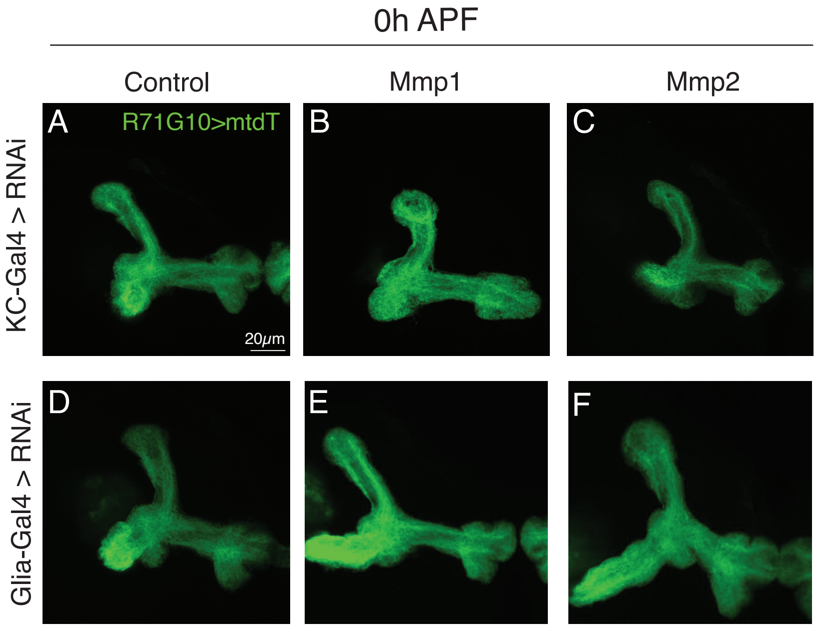
0h APF
Control
Mmp1
Mmp2
KC-Gal4 > RNAi
Glia-Gal4 > RNAi
A
R71G10>mtdT
20μm
B
C
D
E
F

**Figure S7. Glial and neuronal MMPs are not required in KCs nor in glia for initial γ-axon growth**

(A-F) Confocal z-projections of pupal MB axons at 0 hours after puparium formation (h APF). γ-KCs are labeled by R71G10-QF2-driven QUAS-mtdT-HA (green). The indicated UAS-RNAi constructs were driven by the pan-KC driver OK107-Gal4 (A-C), or by the pan-glial driver Repo-Gal4 (D-F). Control (A, D) is RNAi targeting Luciferase. Scale bar corresponds to 20μm.
