## Supplementary material for "A *Drosophila* screen of schizophrenia-related genes highlights the requirement of neural and glial matrix metalloproteinases for neuronal remodeling": Fig. S8

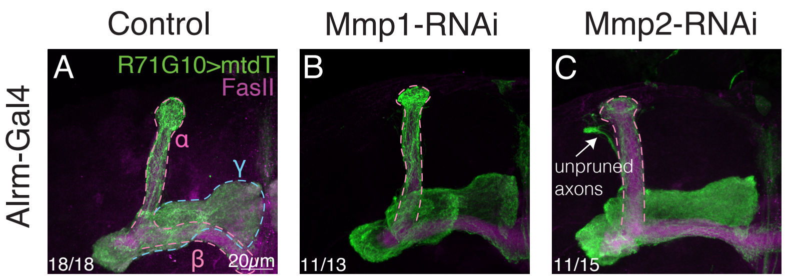
Control
Mmp1-RNAi
Mmp2-RNAi
Alrm-Gal4
A
R71G10>mtdT
FasII
α
γ
β
20μm
18/18
B
11/13
C
unpruned axons
11/15

**Figure S8. Astrocytic Mmp2 is required for γ-axon pruning**

(A-C) Confocal z-projections of adult MBs in which γ-KCs are labeled by R71G10-QF2-driven QUAS-mtdT (green), and control (A; y,w), Mmp1-RNAi (B) or Mmp2-RNAi (C) are expressed in Astrocytes-like glia using Alrm.

The γ lobe is highlighted by a light blue dashed line, and the α/β lobes are highlighted by a magenta dashed line.

FasII antibody (magenta) strongly labels α/β-KCs and weakly labels γ-KCs.

Arrows indicate unpruned axons. Scale bar corresponds to 20μm.
