## Supplementary material for "A *Drosophila* screen of schizophrenia-related genes highlights the requirement of neural and glial matrix metalloproteinases for neuronal remodeling": Fig. S9

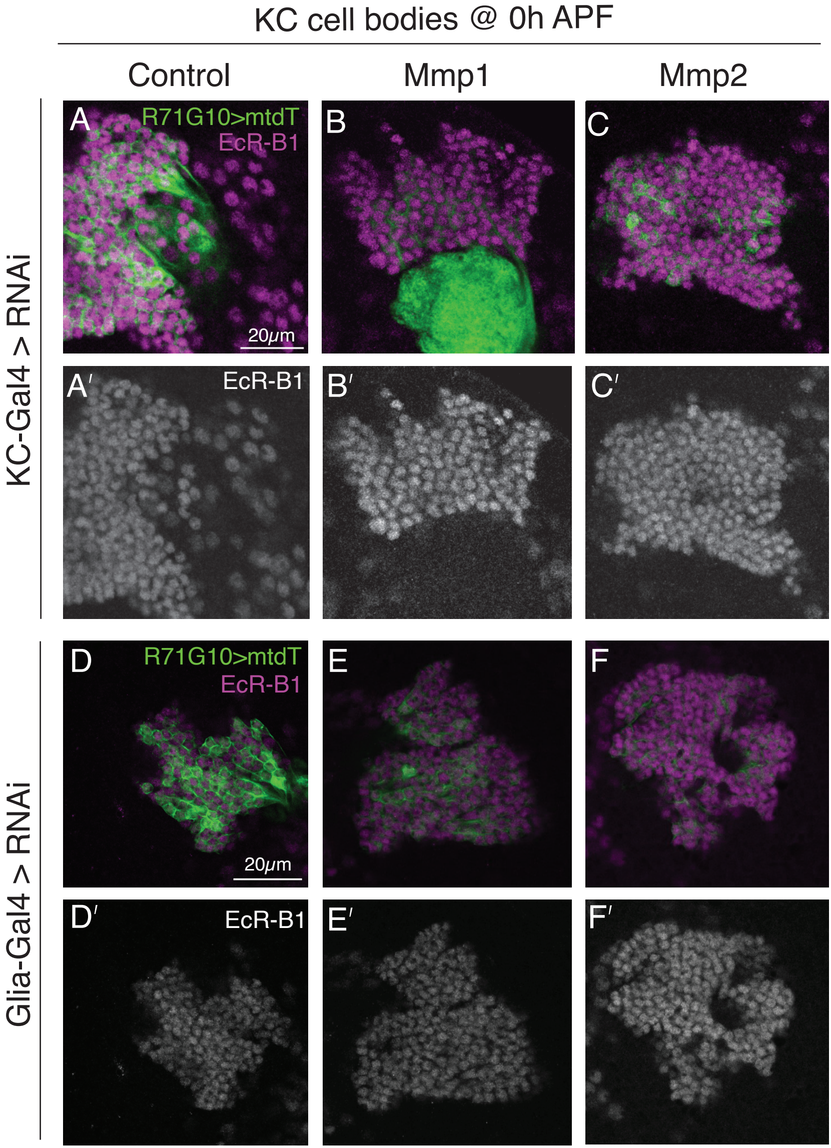
KC cell bodies @ 0h APF
Control
Mmp1
Mmp2
KC-Gal4 > RNAi
Glia-Gal4 > RNAi
A
R71G10>mtdT
EcR-B1
20μm
B
C
A′
EcR-B1
B′
C′
D
R71G10>mtdT
EcR-B1
20μm
E
F
D′
EcR-B1
E′
F′

**Figure S9. MMPs do not act upstream of EcR-B1 in γ-KCs.**

(A-F) Confocal single slices of KC cell bodies at 0 hours after puparium formation (0h APF). γ-KCs are labeled by R71G10-QF2 driven QUAS-mtdT.HA (green), and stained with anti-EcR-B1 antibody (magenta or grey in prime panels). The indicated UAS-RNAi transgenes were driven by the pan-KC driver OK107-Gal4 (A-C) or by the pan-glial driver Repo-Gal4 (D-F). Scale bar corresponds to 20μm.

RNAi transgenes are as follow: Luciferase-RNAi (A, D), Mmp1-RNAi (B, E), Mmp2-RNAi (C, F).
